# Traditional phylogenetic models fail to account for variations in the effective population size

**DOI:** 10.1101/2022.09.26.509598

**Authors:** Rui Borges, Ioanna Kotari, Juraj Bergman, Madeline A. Chase, Carina F. Mugal, Carolin Kosiol

## Abstract

A substitution represents the emergence and fixation of an allele in a population or species and is the fundamental event from which phylogenetic models of sequence evolution are devised. Because of the increasing availability of genomic sequences, we are now able to take advantage of intraspecific variability when reconstructing the tree of life. As a result, substitutions can be more realistically modeled as the product of mutation, selection, and genetic drift. However, it is still unclear whether this increased complexity affects our measures of evolutionary times and rates. This study seeks to answer this question by contrasting the traditional substitution model with a population genetic equivalent using data from 4385 individuals distributed across 179 populations and representing 17 species of animals, plants, and fungi. We found that when the population genetics dynamic is modeled via the substitution rates, the evolutionary times and rates of the two models are well correlated, suggesting that the phylogenetic model is able to capture the time and pace of its population counterpart. However, a closer inspection of this result showed that the traditional models largely ignore the effect of the effective population size, even when it is explicitly accounted for in the substitution rates. Our findings suggest that superimposing population-genetics results on the substitution rates is an effective strategy to study mutation and selection biases, while other data sources (e.g., life history traits or polymorphisms) may need to be additionally integrated to make the traditional substitution models sensitive to the impact of genetic drift. When combined with the known effect of ancestral population size on generating phylogenomic incongruence due to incomplete lineage sorting, our findings provide further evidence that unaccounted-for variations in the effective population size may be one of the primary causes of errors in phylogenetic analyses at shorter time scales.

## Introduction

Models of sequence evolution have a long tradition in phylogenetics. The JC (Jukes and Cantor, 1969), HKY (Hasegawa et al., 1985), and GTR (Tavaré, 1986) are just a few examples of substitution models that are widely used in phylogenetic applications. These models describe the evolution of nucleotides, codons, or amino acids over time, using the substitution process as their baseline event. A substitution represents the sequence of events by which the allelic content of a specific locus in a population is completely replaced. As a result, substitution models are generally quite vague in terms of the processes they may represent but quite flexible, as they implicitly account for any mutation, selection, or fixation biases leading to the fixation of any new allele.

Substitution models were built to characterize fixed differences between species and thus only consider interspecific variation. However, due to remarkable advances in sequencing technologies in recent years, we now have access to genomic data across a wide range of species that allow for the characterization of intraspecific diversity, i.e., polymorphisms. This type of variation is still suboptimally integrated into phylogenetic analyses: e.g., polymorphisms are sometimes treated as sequence ambiguity or error at the tip nodes or totally ignored via consensus sequences. Nevertheless, polymorphisms, which lie at the core of population genetics, have proven to be highly informative for tree reconstruction (Leaché and Oaks, 2017). A classic example is the contribution of polymorphisms in correcting species trees for the effect of incomplete lineage sorting (i.e., the persistence of polymorphisms over several speciation events) (Maddison and Knowles, 2006).

The attempt to incorporate inter- and intra-specific variation into phylogenetic inference resulted in the development of models that, while describing the substitution process, aim to more generally describe how genetic variation changes over time. To do so, they employ a mechanistic description of well-known population forces such as population size variations, mutation rate, selective pressures, and so forth, typically based on the Moran (Moran, 1958) or Wright-Fisher (Fisher, 1930; Wright, 1931) models or the Kingman’s coalescent (Kingman, 1982). Examples of such approaches are the multispecies coalescent (Rannala and Yang, 2003) and the polymorphism-aware phylogenetic models [PoMos; De Maio et al. (2013, 2015)]. Guided by empirical studies, these models are becoming more realistic, accounting for other essential processes such as migration (Flouri et al., 2020) or nucleotide usage biases (Borges et al., 2019b).

As more genomic sequences become accessible, it will soon be possible to account for polymorphisms and the processes governing their dynamic when determining the evolutionary history of more recent groups of the tree of life. In this process, many clades and divergence times will undoubtedly be revisited and revised. While it is obvious that different evolutionary models provide different interpretations of evolutionary times and rates, it is still unclear whether population-genetics and phylogenetic narratives are consistent in general. Determining the evolutionary significance of population processes playing a role in sequence evolution will further help us to understand which are the ones that are likely to be neglected and which are already well captured by the traditional approaches.

Here, we used population genomic data from abundantly sequenced biologically diverged taxa across animals, plants, and fungi to compare the traditional substitution model with a population-genetics counterpart that accounts for mutation bias, selection, and genetic drift. Specifically, we compared these models in terms of divergence rates and substitution times. This study aims to determine the evolutionary contexts in which these models may exhibit different time and pace and to identify the evolutionary processes responsible for such discrepancies.

## Results and Discussion

### Modeling the substitution process at two different levels of detail

The substitution process can be modeled at various levels of complexity. In this study, we will concentrate on two different models: a simpler one inspired by phylogenetic models of sequence evolution (which we will refer to as *M*_1_ or substitution model) and a more complex one that takes into account some population-level forces that are likely involved in the substitution process (which we will refer to as *M*_2_ or Moran model). In both cases, the substitution process is modeled as a Markov chain that describes the evolutionary dynamic between two alleles that are present at the same locus in a population or species. A valuable property of both models is that we can easily derive closed-form expressions for the stationary distribution, divergence rate, and substitution time, quantities that we here used to compare these models. We note that substitution models are usually modeled as a continuous-time process, whereas here we used a discrete-time variant to be comparable to the Moran model. However, these models are mathematically equivalent for the aforementioned quantities (text S1), permitting us to extend our comparisons to the continuous case.

Let us consider a model *M*_1_ with only two states, *S* and *W*, and describing the substitution process with only two rates: λ_*WS*_ is the probability of a *W* → *S* change per time unit, and *vice versa* for λ_*SW*_ (figure 1A). The model *M*_2_ describes the change in the frequencies of *S* and *W* over time in a population of size *N* (figure 1B). In this model, we consider the existence of fixed states, in which all *N* individuals have the same allele, as well as polymorphic states, in which the alleles *S* and *W* coexist at different frequencies. To describe the substitution process in this population, we use a Moran model with boundary mutations and selection.

**Figure 1:**
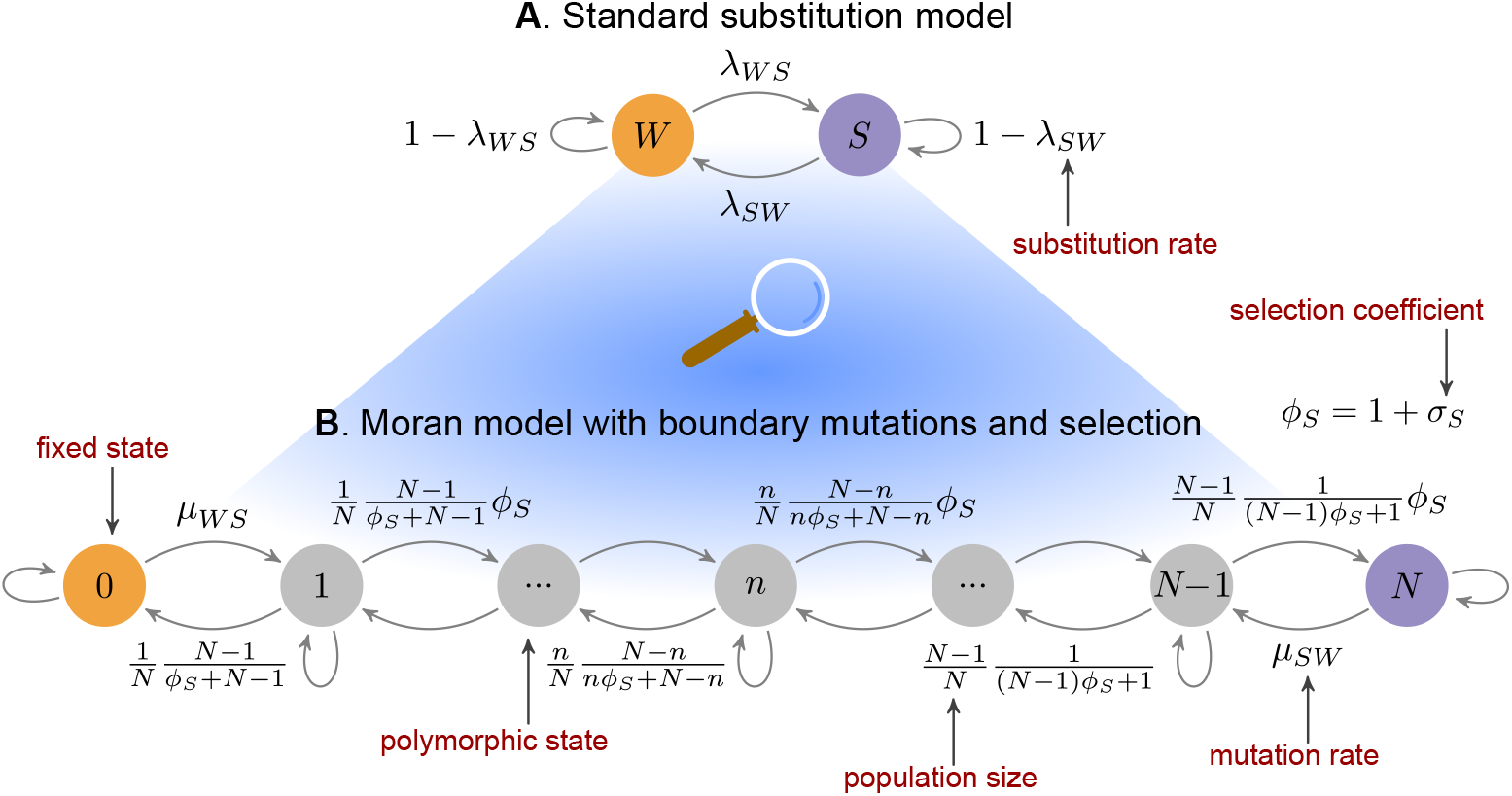
Two possible models for the substitution process. **(A)** A simple substitution model inspired by the phylogenetic models of sequence evolution. The two alleles *W* and *S* interchange each other with substitution probabilities λ_*WS*_ and λ_*SW*_. **(B)** A Moran model with boundary mutations and selection. The states of this model represent the frequency of the *S* allele in a population of *N* individuals: 0 and *N* represent fixed states, while the remaining ones are polymorphic. The probabilities of a mutation from *W* to *S* and *S* to *W* are respectively denoted by *µ*_*WS*_ and *µ*_*SW*_. Under the Moran model, the probability of choosing one of the alleles to die and another to reproduce is 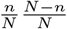. In the figure, this ratio is expanded to account for directional selection, where *σ*_*S*_ is the selection coefficient of the *S* allele; we assume *σ*_*W*_ = 0. For the sake of clarity, the probabilities of remaining in the same state were not depicted.

The probabilities of a mutation per Moran event from *W* to *S* and from *S* to *W* are represented by *µ*_*WS*_ and *µ*_*SW*_, respectively. Mutation rates are assumed to be boundary (or low), biased, and reversible: low, so that by the time a new mutation occurs, the previous one has been fixed [this is a reasonable assumption for most eukaryotes (Lynch et al., 2016)]; biased, because *µ*_*WS*_ and *µ*_*SW*_ may differ; reversible, because multiple mutations per site are permitted. The Moran model models genetic drift by assuming that one random individual is chosen to reproduce and one random individual is chosen to die at each time step (Moran, 1958). These probabilities depend on the population size *N*, which determines the strength of genetic drift, and the selection coefficient *σ*_*S*_, which confers a reproductive advantage to individuals carrying an allele *S. σ*_*S*_ relates to the fitness coefficient via the equation *ϕ*_*S*_ = 1 + *σ*_*S*_. Four-allelic versions of this model have been previously used for phylogenetic analyses (Borges et al., 2019a; Borges and Kosiol, 2020; Borges et al., 2022).

Although models *M*_1_ and *M*_2_ allow for the study of the substitution process, they do so in different ways. There are, however, certain considerations that can be made in order to compare them. If a single individual is sampled from a population, that individual can carry either a *S* or *W* nucleotide per genomic position, but never a polymorphism. This sampling takes the *M*_2_ model back to the state space where the substitution model operates, thus representing a natural approach to establishing a relationship between these models. The probabilities *p*_*W*_ and *p*_*S*_ of sampling a single individual carrying a *W* or a *S* allele can be calculated if we assume that the population is at or close to the stationary state:

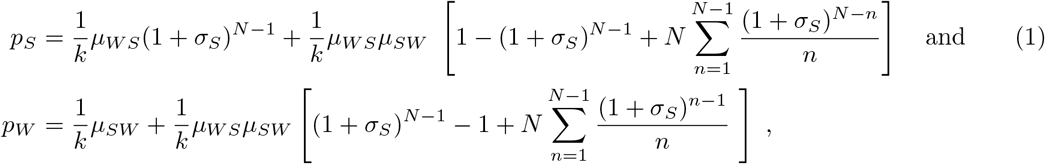

where *k* is a normalization constant [equation S10 in text S2]. We expect these probabilitites to match the stationary frequencies of the states *W* and *S* in the *M*_1_ model, as both represent observed fixed sites. By relating these quantities (text S3), we obtain that the substitution rates can be expressed by

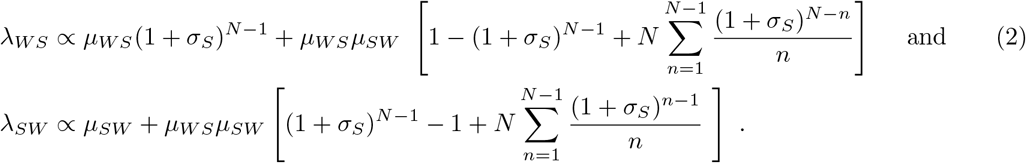

The relation depicted in the equation (2) is just another example of the use of population genetics-informed parameters in phylogenetic models. Similar strategies have been used by several authors, such as Halpern and Bruno (1998); Robinson et al. (2003) in developing the first mutation-selection codon models or Lartillot (2013) to study the dynamic of GC-biased gene conversion (gBGC) across mammals. In the following sections, we used identity (2) to redefine the substitution rates of model *M*_1_, making it thus dependent on the same evolutionary processes that govern the *M*_2_ dynamic.

### The evolutionary dynamic between strong and weak nucleotides

To make our model comparisons more biologically insightful, we used population parameters estimated from empirical data. We concentrate on the dynamic of strong (*S*: *G* or *C*) and weak (*W* : *A* or *T*) nucleotides. One of the reasons for selecting such a system is that it is observed that the composition of strong and weak nucleotides is regulated throughout the genome by two antagonistic processes: while mutations are generally found to be biased towards weak nucleotides (Keightley et al., 2009; Lynch, 2010), another molecular process, known as gBGC, acts in the opposite direction. gBGC is a recombination-associated process that induces a transmission distortion towards strong nucleotides at *WS* heterozygote sites, which ultimately promotes the fixation of strong alleles (Duret and Galtier, 2009). Furthermore, this nucleotide dynamic is thought to act on a genome-wide scale and has been observed across a wide range of taxa (Pessia et al., 2012; Galtier et al., 2018). Due to the fact that the substitution rates represent the combined effect of these antagonistic processes (along with the effective population size that controls their efficiency), the weak-strong system is exceptionally well suited for testing the effects of the population-genetics dynamic in the substitution models.

Under the weak-strong allelic system, *µ*_*WS*_ and *µ*_*SW*_ are the mutation rates from weak to strong and strong to weak alleles, respectively. gBGC is typically modeled equivalent to directional selection (Nagylaki, 1983). *σ*_*S*_ is the selection coefficient of strong nucleotides or the GC-bias preference. These parameters, along with the effective population size *N*, were estimated using allele counts from various genomic positions, assuming that a sample of *M* individuals was drawn from a population of *N* individuals with a mutation-selection dynamic close to or at the steady-state (text S3). The inferences were conducted using a Bayesian framework, which estimates the population parameters described above using empirically obtained mutation rates. We created a program called BESS (Bayesian Estimator of the Site-frequency Spectrum) that performs the aforementioned steps. More information on BESS and the inferential analyses can be found in the Materials and Methods section.

Our data set includes 179 populations from six different taxa, including great apes, mice, flycatchers, fruit flies, Arabidopsis, and baker’s yeast (tables 1 and S1). These taxa were chosen because genome-wide data are available and they are particularly well-characterized in terms of their intraspecific variation. An average of 145 million genomic sites (ranging from 150 thousand to 26 million sites) were used for the inferences, with approximately 26 (ranging from 2 to 243) individuals sampled per population. It was our intention to cover a wide range of population dynamics with variable population sizes, mutation rates, and GC preferences. Figure 2 shows the estimated population parameters per population, and how they vary significantly within and between taxa. To better assess the magnitude of these forces, the mutation rates and the selection coefficients are scaled with the population size.

**Table 1:** Population genomic data used in this study.

| Taxa | Species and no. of populations | Type of genomic sites and source | Mutation rate and source |
| --- | --- | --- | --- |
| Great apes | *Pongo pygmaeus* (1), *P. abelii* (1), *Gorilla gorilla* (3), *Pan troglodytes* (4), *P. paniscus* (1) and *Homo sapiens* (1) | Four-fold degenerate sites (Bergman and Schierup, 2021) | $25 \times 10^{-9}$ (Venn et al., 2014) |
| Mice | *Mus musculus* (8) and *Mus spretus* (1) | Whole genome (Harr et al., 2016) | $5.4 \times 10^{-9}$ (Uchimura et al., 2015) |
| Flycatchers | *Ficedula albicollis* (1), *F. hypoleuca* (1), *F. parva* (1), *F. albicilla*, *F. speculigera* and *F. semitorquata* (1) | Four-fold-degenerate sites (Burri et al., 2015; Chase and Mugal, 2022) | $4.6 \times 10^{-9}$ (Smeds et al., 2016) |
| Fruit flies | *Drosophila melanogaster* (30) | whole genome (Hervas et al., 2017) | $3.5 \times 10^{-9}$ (Keightley et al., 2009) |
| Arabidopsis | *Arabidopsis thaliana* (37) | coding regions (Initiative, 2000) | $7.0 \times 10^{-9}$ (Ossowski et al., 2010) |
| Baker's yeast | *Saccharomyces cerevisiae* (86) | whole genome (Peter et al., 2018) | $0.3 \times 10^{-9}$ (Lynch et al., 2008) |

**Figure 2:**
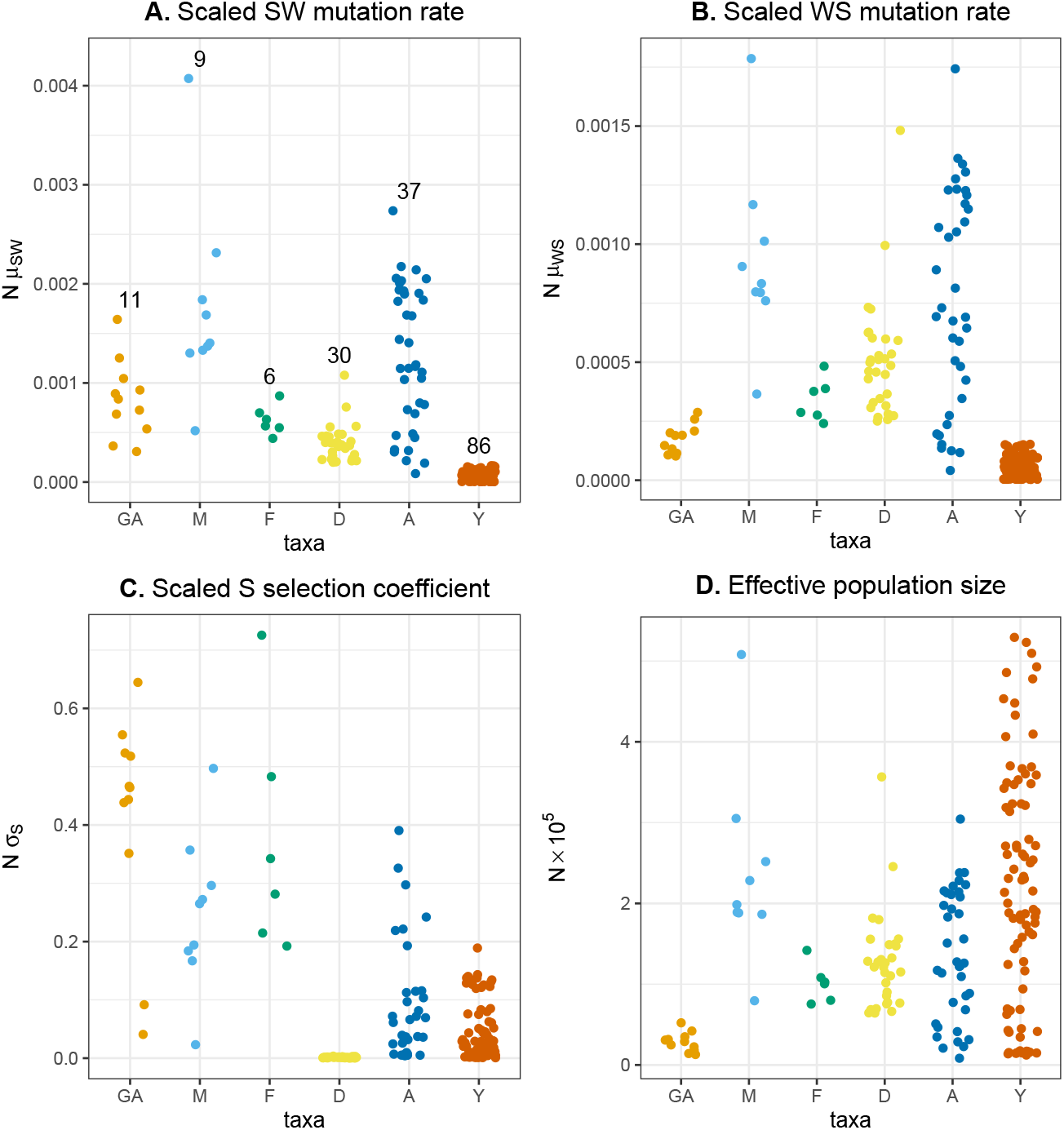
Population estimates of the forces at work in the weak vs. strong allele dynamic. To better assess the magnitude of these forces, the mutation rates and selection coefficients were scaled with population size. The parameter estimates were obtained with BESS (see Materials and Methods for more details) using allelic counts from population genomic data. The numbers depicted in panel A represent the number of populations per taxon. Legend: great apes (GA), mice (M), flycatchers (F), fruit flies (D), arabidopsis (A), and baker’s yeast (Y).

Strong to weak base mutations were found to be more frequent than weak to strong base mutations, which is consistent with other studies (Keightley et al., 2009; Lynch, 2010) (figure 2A–B). We also observed that the selection coefficients of strong bases varied greatly between taxa and populations, being quite pronounced in great apes, mice, and flycatchers but almost nonexistent in fruit flies (figure 2C). Our results are supported by studies showing that the composition of weak and strong bases is driven by gBGC in great apes (Borges et al., 2019a), mice (Galtier, 2021), flycatchers (Bolívar et al., 2016), Arabidopsis (Günther et al., 2013) and baker’s yeast (Lesecque et al., 2013), but not in fruit flies (Robinson et al., 2014). However, our results also showed that selection in favor of strong bases varies considerably among taxa and correlates negatively with the effective population size (*ρ* = 0.70 *p*–value*<*0.001; figure S3). Since the recombination landscape is remarkably conserved between closely-related populations and species, we attribute this variation to nonequilibrium conditions related to fluctuating demography or gBGC (we note that our model assumes that both processes are constant) (Brandvain and Wright, 2016; Müller et al., 2022). In any case, as observed in other studies (Glémin et al., 2015; Bolívar et al., 2019), the estimated GC preferences appear to operate in the nearly neutral range for all taxa (i.e., *Nσ*_*S*_ *<* 1).

The population size varies significantly among taxa, ranging from around 4 500 individuals in great apes to about 530 000 in yeasts (figure 2D). Other studies have estimated far larger population sizes for fruit flies and yeast in the order of millions of individuals [e.g., Hodgins-Davis et al. (2015)]. These differences may be attributed to different model assumptions. Concretely, we account for reversible and biased mutations as well as fixed sites, unlike many estimators that assume the infinite sites model and rely solely on polymorphic sites. Fixed sites are by far the most common type among the genomic positions sampled, and we expect them to dominate our estimates of mutation rates, selection coefficients, and effective population sizes. We further note that our estimates of mutation rates, selection coefficients, and population sizes vary independently of the different types of genomic positions used to perform the inferences (whole genome *vs*. coding sequences; table 1).

### Comparing the pace and time of traditional and population-genetics models

Evolutionary rates and times represent important aspects of evolutionary studies and are extensively used in comparative genomics to establish organismal relationships and estimate divergence times. We compared the models *M*_1_ and *M*_2_ in a rate and time dimension by making use of respectively the expected divergence rate (or the expected number of state changes per unit of time) and substitution time from a weak to a strong base.

The divergence rate in the substitution model *M*_1_ simply expresses the expected number of substitutions per unit of time. Differently, in the Moran model *M*_2_, it represents the expected number of mutations and frequency shifts per Moran generation (or *N* Moran events). These quantities can be obtained formally (text S1 and S2) if we assume that the population is at or near to the steady state:

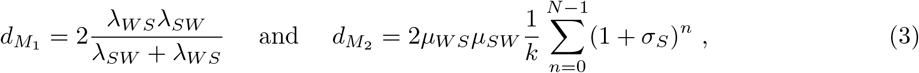

We used the substitution time from a weak to a strong base to establish a time dimension comparison between the models. Despite representing the same quantity, these are expressed differently by the two models. The weak to strong base substitution time in model *M*_1_ represents a single step from a *W* to a *S* state, whereas in the *M*_2_ model is the time it takes to reach the boundary state *N* from the boundary state 0, which includes the time to a mutation plus the fixation time. Both models have closed-form expressions for the substitution time:

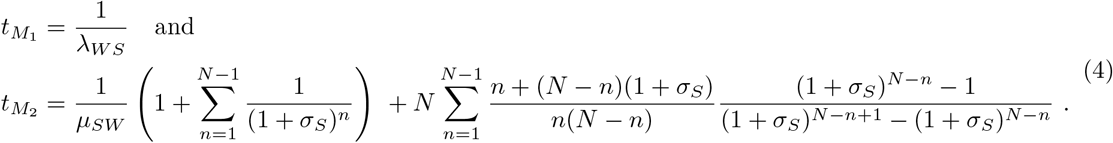

We further replaced the substitution rates λ_*WS*_ and λ_*SW*_ using the relation (2), so these rates and times are comparable and explicitly dependent on the same population processes.

In contrast to the Moran model, which operates in Moran events or generations, the time units of the substitution model are unknown, as we cannot identify the substitution rates based solely on the frequency of *W* and *S* bases. This is also the reason why phylogenetic distances are typically normalized, so that one substitution per unit of time is expected. The implications of this unidentifiability of the substitution rates are that the absolute values of the evolutionary rates and times of the substitution model cannot be interpreted directly. However, as we are more interested in the degree to which they correlate with their population counterparts or their relative change, this aspect does not represent a limitation for our comparisons.

To assess how both models measure evolutionary rates and times, we correlated the normalized divergence rates and substitution times of the two models using the population parameters inferred with BESS and the identities (3) and (4). We observed that the divergence rates are significantly correlated (correlation coefficient *ρ*_*B*_ = 0.733, *p*_0_ *>* 13.815; figure 3A and S6A). *ρ*_*B*_ is corrected for the heterogeneous number of populations per taxa; significant correlations are obtained for *p*_0_ *>* 2.944 (see Materials and Methods for more details on the bootstrap analyses). Divergence rates in the *M*_2_ model, on the other hand, vary significantly more across populations than in the *M*_1_ model. For example, the divergence rates of fruit flies are almost constant under model *M*_1_, whereas they vary considerably in model *M*_2_. Similarly, the substitution times measured by the two models are significantly and strongly correlated (*ρ*_*B*_ = 0.845, *p*_0_ *>* 13.815; figure 3B and S6B) and that the *M*_2_ model captures considerably more variation at substitution times.

**Figure 3:**
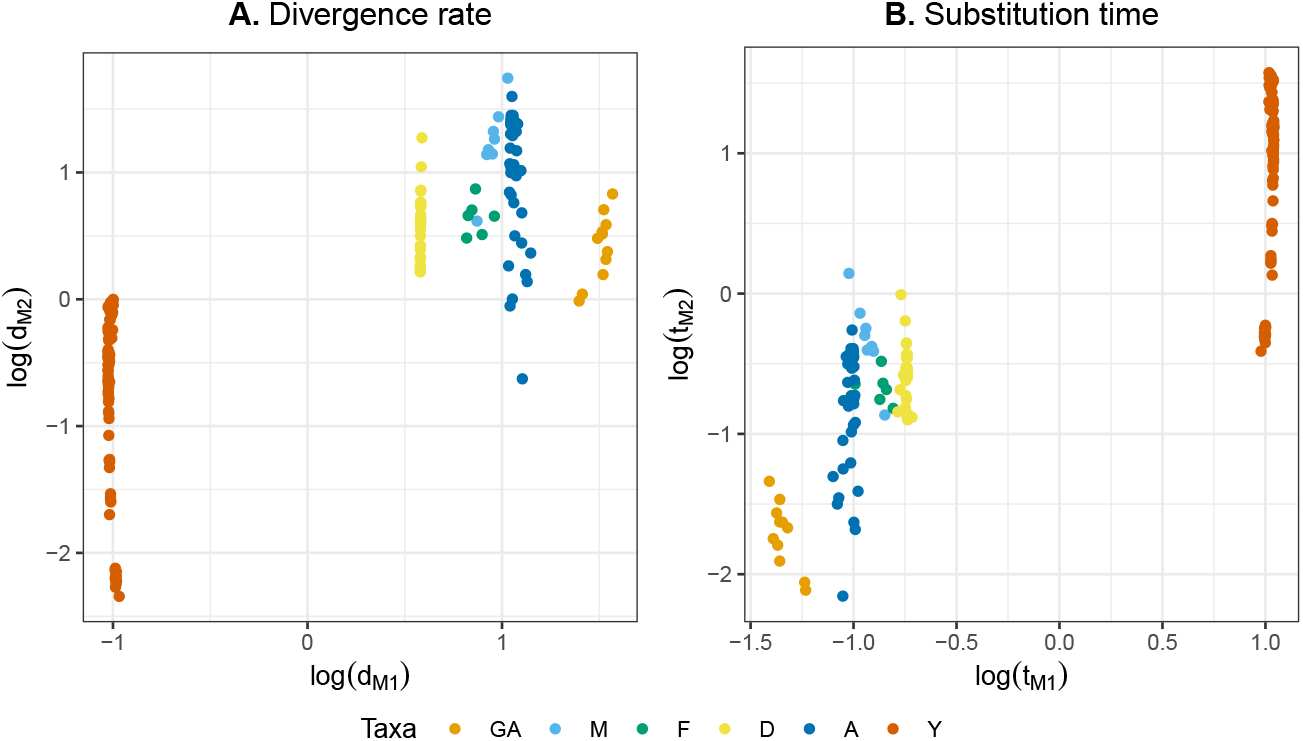
Comparing divergence rates and substitution times between traditional substitution and the Moran model. Comparison between the **(A)** log divergence rates and **(B)** log substitution times in the standard substitution model *M*_1_ and the Moran model with boundary mutations and selection *M*_2_. Legend: great apes (GA), mice (M), flycatchers (F), fruit flies (D), arabidopsis (A), and baker’s yeast (Y).

To better understand the lower variability in the *M*_1_ estimates, we conducted a sensitivity analysis. More specifically, we investigated the effect of different population forces on the estimates of divergence rates and substitution times under the two models. To do that, we perturbed the estimates of each population parameter at a time by *±*10% and calculated the relative difference between the perturbed and unperturbed rates and times. The expectation was that the higher the relative difference, the more sensitive the model is to variations in the population parameter that was perturbed. Figure 4 shows the relative difference in divergence rates and substitution times for each population parameter and model.

**Figure 4:**
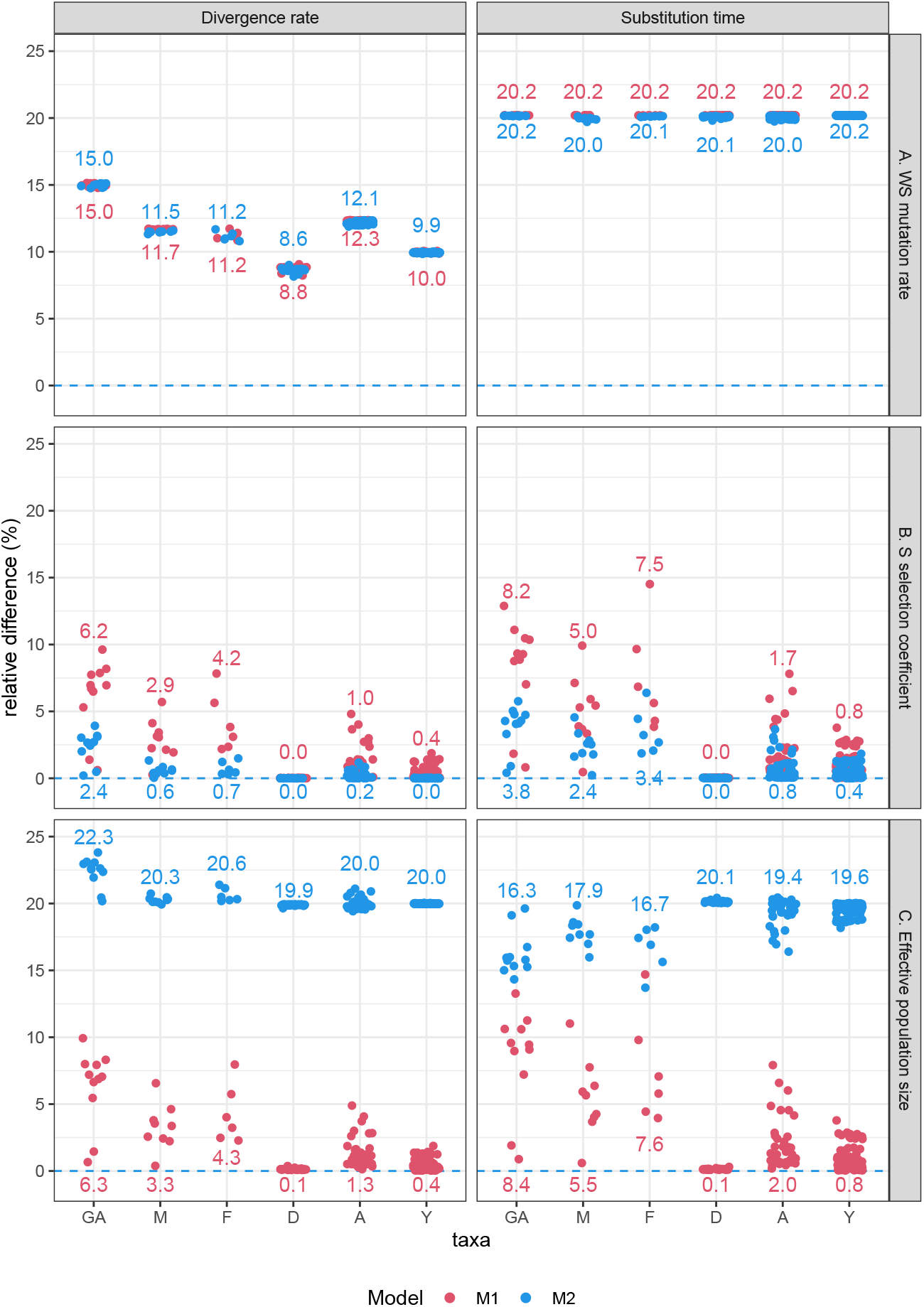
Sensitivity analyses for the divergence rate and substitution time. Left panel: Sensitivity analyses for the divergence rate. The sensitivity analyses for the *SW* mutation rate can be found in figure S4. Right panel: Sensitivity analyses for the substitution time. Because we are only considering the *WS* substitution time, the *µ*_*WS*_ has no measurable effect [equation (4)]. The relative difference expresses the effect of changing each population parameter by *±*10% on the divergence rates. The numbers represent the averaged relative divergence across all populations. Legend: great apes (GA), mice (M), flycatchers (F), fruit flies (D), arabidopsis (A), and baker’s yeast (Y).

The sensitivity analyses showed that the impact of mutation rates is overall consistent between the two models: 5 – 15% for the case of the divergence rate and around 20% for the substitution time (figure 4A and S4), suggesting that their impact is likely to affect inference in both models.

The GC-bias rate, on the other hand, has a smaller effect: 0 – 6.2% for the case of the divergence rate and 0.0–8.2% for the substitution time; figures 4B). An intriguing feature is that the impact of selection on the divergence rate and substitution is on average greater in the *M*_1_ model than in the *M*_2_ model (figures 4B). This finding suggests that using population genetics-informed substitution rates may slightly overestimate the contribution of nucleotide usage biases to the overall evolutionary process. Our findings also showed that the impact of selection is clearly more pronounced in taxa where GC-bias is stronger, such as the great apes, mice, and flycatchers, and has almost no effect in taxa where GC-bias is weak, such as baker’s yeast, or is almost absent, as in fruit flies. This suggests that the substitution models may only be able to capture the net effects of selection. This result is expected by the nearly neutral theory, in which the substitution rates might be accelerated or decelerated depending on the population-scaled selection coefficient (Ohta, 1973). In fact, if we expand the selection term of the relations (2) in its Maclaurin series and further assume that *σ*_*S*_ is small, we obtain

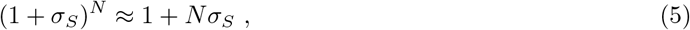

from which we can recapture the scaled selection coefficient. We further note that these results on selection must be generalized with care since here, we only investigated a particular case of directional selection acting on GC alleles. As shown by our results (figure 2C) and other studies [e.g., Galtier et al. (2018); Bolívar et al. (2019); Borges et al. (2022)], although ubiquitous, gBGC is likely to act in the nearly neutral range. Thus, despite our results showing selection has a minor effect on substitution times and rates, one should not underestimate the effect of other selection forces in the substitution process. For example, purifying selection acting on coding regions is known to be both pervasive and strong (Cvijović et al., 2018).

The effective population size follows a completely different pattern than that of the selection coefficients and mutation rates (figure 4C). Changes in population size may explain approximately 20.0 – 22.3% of variation in substitution rates in the *M*_2_ model, but only 0.1– 6.3% percent in the *M*_1_ model (left panel of figure 4C). Similarly, the effective population size is the driving force explaining the main differences in the estimated substitution times between models: while it accounts for 16.3 – 20.1% of its variation in the *M*_2_ model, it only accounts for 0.1 – 8.4% in the *M*_1_ model (right panel of figure 4C). These results suggest that the effect of the effective population size, or genetic drift, is largely ignored in the substitution model. We also observed that populations with stronger GC-bias (great apes, mice and flycatchers) have a more pronounced effect of population size on the substitution rates estimated by the model *M*_1_. This is likely because the population size influences the substitution rates through the selection coefficient [as shown in equation (2)].

We conclude that while evolutionary rates and times correlate well between the traditional substitution model and its population-genetics equivalent, the former considerably underappreciates genetic drift, despite the effective population size being explicitly modeled into the substitution rates. We note that this effect should be even more pronounced, as in our models we assumed a fixed effect of demography, whereas in natural populations, it is likely to vary over time.

### The impact of unseen variation on the substitution process

The previous section established that traditional phylogenetic models tend to disregard the effect of population size in their evolutionary rates and times. This conclusion was drawn based on an analysis of a single individual sample. However, as more genomic sequences become available and we are able to sample more polymorphisms, it is worth investigating how much they are likely to affect our inferences of evolutionary rates and times.

A well-known consequence of sampling alleles from a population of *N* individuals is that one always assesses a subset of the existing diversity. This bias occurs primarily because low-frequency polymorphisms are more likely to be sampled as monomorphic. In equation (1), we established the probability of an *S* and *W* base when a single individual is sampled from the population, but we can generalize this sampling procedure to *M* individuals. More formally, the probability *c*_*M*_ of sampling *M S* alleles from the population is

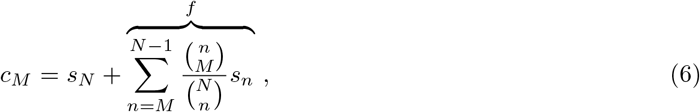

where *s*_*n*_ is the expected stationary frequency of a population having *n S* alleles and (*N* − *n*) *W* alleles. Equation (6) quantifies the contribution of true fixed (first term) and polymorphic sites (second term) to the frequency of observed fixed sites of type *S*. The second term that we denoted by *f* is particularly important because it represents the excess of fixed sites, i.e., the proportion of polymorphic sites that are likely to be sampled as monomorphic in a sample of *M* individuals. *f* becomes less important as *M* approaches *N* ; however, this excess is also affected by the mutation-selection dynamic via the stationary quantity *s*_*n*_. As a result, the relative contribution of the dynamic and sampling effects to the excess of sampled fixed sites is not immediately clear.

To investigate the contribution of sampling polymorphisms to the observed excess of fixed sites, we performed a correlation analysis between the excess of fixed sites and the proportion of polymorphic sites in the data. We found a weak and non-significant correlation (*ρ*_*B*_ = −0.191, *p*_0_ = −0.826; see Figure 5A and S7A). Additionally, we examined the correlation between the excess of fixed sites and the sample size, as a larger sample size is expected to result in a smaller excess of monomorphic sites. However, we again observed a weak and non-significant correlation (*ρ*_*B*_ = 0.077, *p*_0_ = 0.372; see Figure 5B and S7B). Notably, we found that the relationship between these variables varied considerably across taxa, with no clear trends observed when correlations were conducted per taxa; some correlations were positive, others were negative, and not all were significant (see Table S2). The lack of strong correlations between the excess of fixed sites and the number of sampled individuals or polymorphisms suggests that the population dynamic, and not the sample size, may be the primary driver of the observed sampling bias.

**Figure 5:**
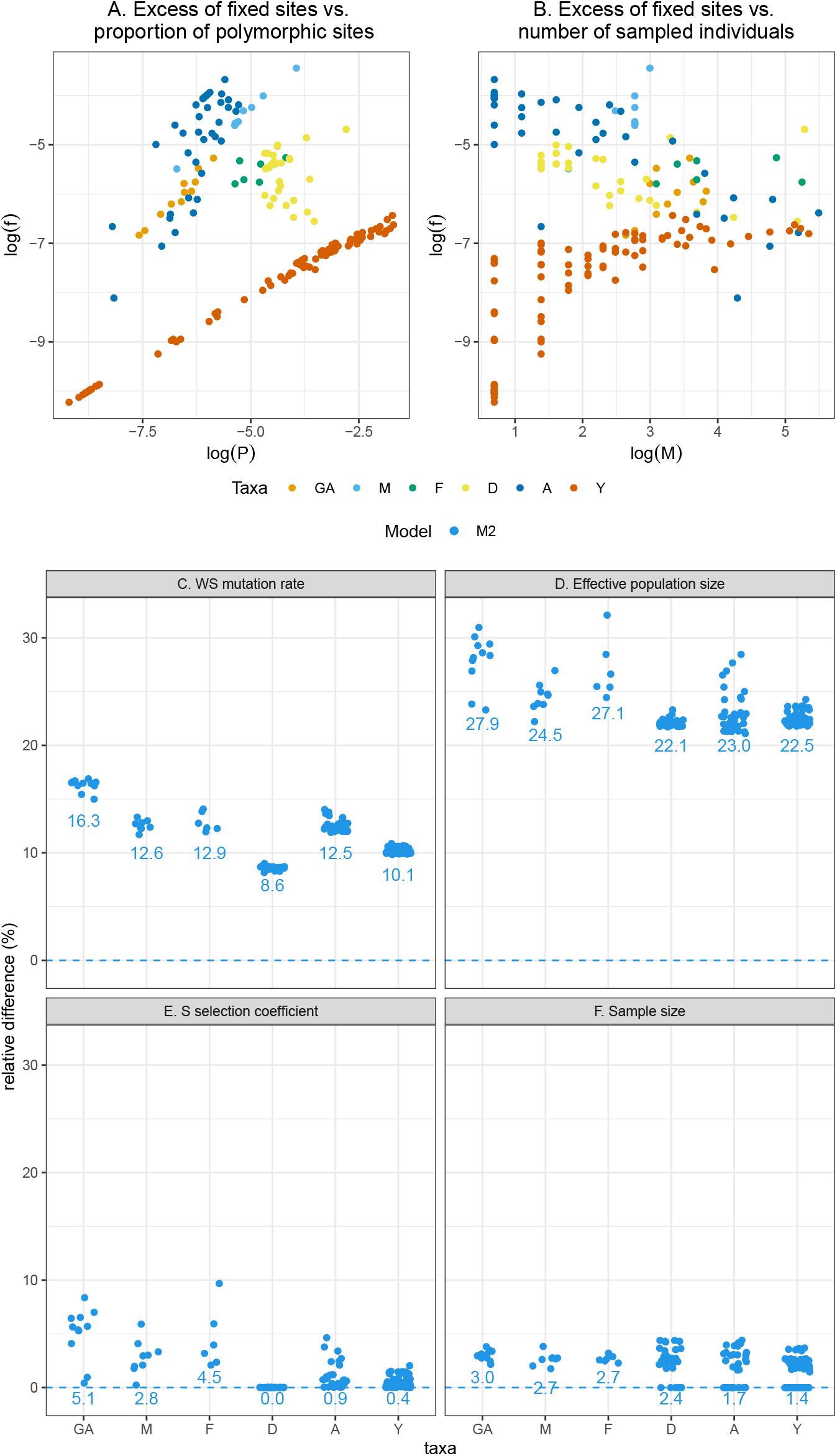
Assessing the relative contribution of the population dynamic and sampling of individuals to the excess of observed fixed sites. **(A–B)** Comparison between the log excess of fixed sites (*f*) and proportion of polymorphism (*P*) and sample size (*M* ; for diploid organisms, *M* represents the number of haploid individuals). **(C–F)** Sensitivity analyses for the different population forces. The relative difference expresses the effect of changing each population parameter by *±*10% on the excess of observed sites [equation (4)]. The plot relative to the impact of *SW* mutation rate can be found in figure S5. The numbers represent the average relative divergence across all populations. Legend: great apes (GA), mice (M), flycatchers (F), fruit flies (D), arabidopsis (A), and baker’s yeast (Y).

**Figure 6:**
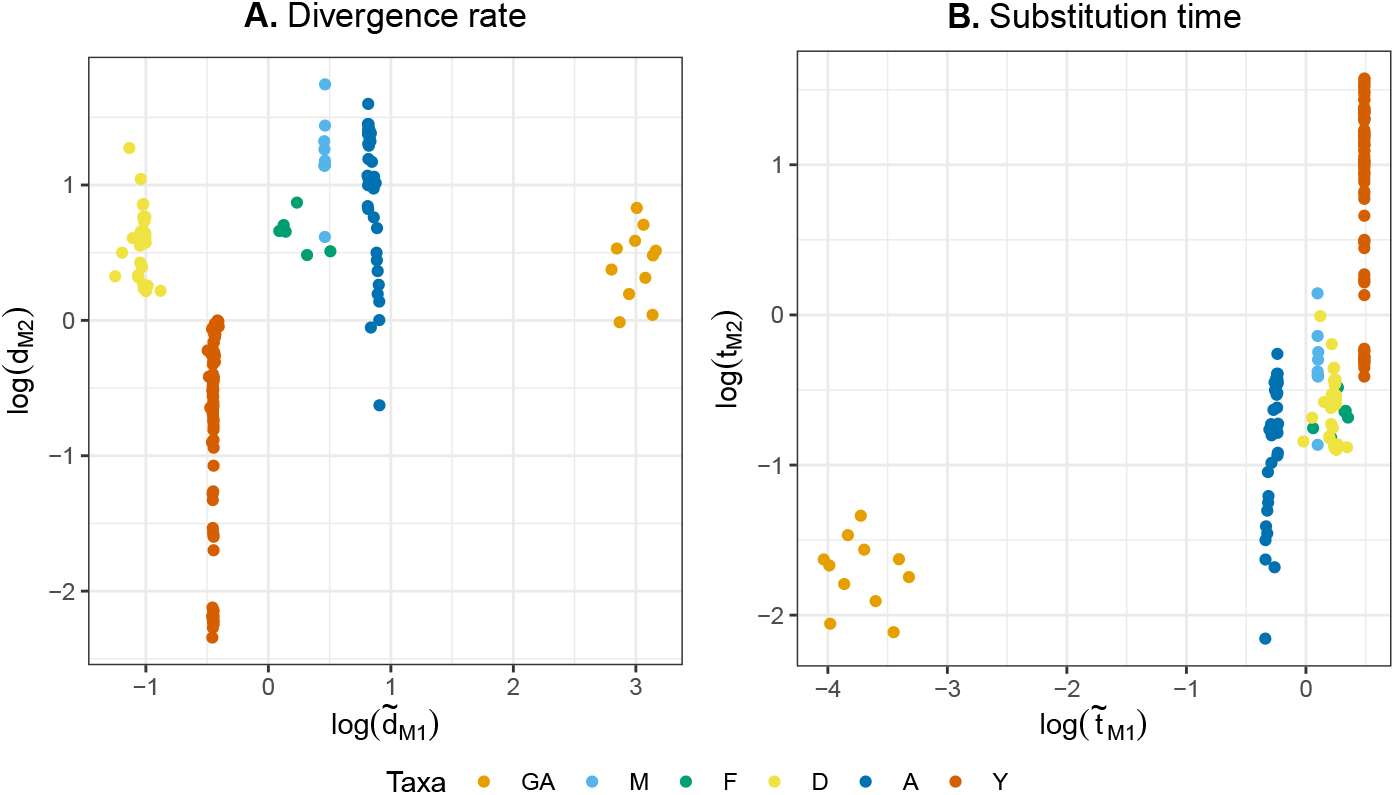
Comparing rates and times between population-informed and population-naïve substitution models. The tilde marks the models for which the substitution rates were directly estimated from the observed frequencies of *W* and *S*. Legend: great apes (GA), mice (M), flycatchers (F), fruit flies (D), arabidopsis (A), and baker’s yeast (Y).

In order to determine the extent to which the population dynamic and sample size contribute to an excess of fixed sites, we conducted a sensitivity analysis. Given that the excess of fixed sites cannot be adequately assessed using the model *M*_1_, which does not take polymorphisms into account, we solely used model *M*_2_ for these analyses. Our findings indicate that the mutation rates and effective population size have the greatest influence on the excess of observed fixed sites, explaining approximately 8.6% to 16.3% (see Figure 5C and S5) and 22.1% to 27.9% (see Figure 5D), respectively. The selection coefficient and sample size had a minor impact, accounting for approximately 0.0% to 5.1% (see Figure 5E) and 1.4% to 3.0% (see Figure 5F), respectively.

It is evident that the sample size has a relatively minor impact on sampling biases compared to the population dynamic. We can therefore conclude that modeling the mutation-selection-drift dynamic may not only aid in correcting for the excess of fixed sites (thus resulting in less biased assessments of the allele composition), but also result in more accurate assessments of evolutionary times and rates. This is observation is supported by the significant correlations observed for the divergence rates and substitution time (figure 3), where the population dynamic is modeled via the substituion rates. It should be noted that the number of individuals sampled is just one component of our sampling process, with the number of genomic sites being another factor. As acquiring a larger number of sites is not currently a sampling challenge, we did not address it here. We further note that despite differences in the number of analyzed sites (9 million for great apes, 26 million for mice, and 1 million for flycatchers; see table S1), all three species exhibit similar patterns with regard to the impact of selection and population size.

Thus far, we have represented the substitution rates of model *M*_1_ using a population-genetic description of the substitution process [equation (2)]. Yet, in most phylogenetic applications, there is no information about the underlying population dynamic or genomic data that allows one to infer it. Consequently, phylogeneticists assume that the observed sites are all fixed. However, according to our results, this assumption may be problematic, as the population dynamic seems to be fundamental in accurately determining the excess of fixed sites. To address this, we re-estimated the divergence rate and substitution time of model *M*_1_ directly from the empirical frequencies of *W* and *S* bases. This enabled us to test how evolutionary rates and times change when they are inferred directly from the observed fixed sites. As a result of this re-estimation, weaker and, in the case of the divergence rate, insignificant correlations were observed: *ρ*_*B*_ = 0.173 (*p*_0_ = 1.631; see 6A and S8A) and *ρ*_*B*_ = 0.733 (*p*_0_ *>* 13.815; see 6B and S8A).

These correlations suggest that when the substitution rates are estimated directly from the observed fixed sites, they might not express the tempo and pace of the population process. This result is somewhat predictable because fixed sites are the result of a tug-of-war between mutation, selection, and genetic drift, and their impact on the evolutionary process should be highly sensitive to biased estimates of fixed site frequencies.

## Conclusion

In this study, we used population genomic data from diverse taxa across animals, plants, and fungi to compare the evolutionary rates and times between the traditional phylogenetic model and a comparable population-genetics model. Our approach was to modify the substitution rates by incorporating a population dynamic that takes into account mutation, selection, and genetic drift. Our findings suggest that adopting this approach enables the substitution models to reproduce the timing and rate of the underlying population genetics model. However, we also found that the impact of genetic drift is largely underappreciated. These findings have three important implications for phylogenetic analyses and inference, particularly at shorter time scales.

First, our results demonstrate that the incorporation of population-informed formulae into substitution rates is an effective method for analyzing the influence of mutation and selection bias on the evolutionary process. This indicates that reducing the evolutionary process to the substitution event is still sufficient for investigating the role of mutation and selection in determining the genome’s base composition. This result is also supported by a recent study by Latrille et al. (2022), which demonstrates that phylogenetic codon models and population-genetic tests of adaptation are congruent.

Second, we found that when estimating evolutionary rates and distances using the empirical base composition, their time and pace might differ from their population counterpart. This suggests that if polymorphisms or other information on the population dynamic is available, it should be used to correct the base composition frequencies at the tip nodes, as opposed to less optimal approaches such as the use of consensus sequences or tip corrections that handle ambiguity and error but are not specially tailored for the sampling of individuals from a population. Corrections for this sampling of individuals based on the binomial distribution have been proposed in other phylogenetic contexts [e.g., Schrempf et al. (2016); De Maio et al. (2015)], but can be easily adapted to the traditional models. Here, we utilized exceptionally well-characterized taxa, for which an abundance of polymorphic data exists. Unfortunately, it may not always be possible to account for bias in the base composition, as this information does not exist for nonmodel organisms.

Third, our findings show that the traditional models may be incapable of establishing meaningful distances between closely related taxa. We could attribute this pattern to a nonidentifiable effect of the effective population size on the evolutionary times and rates, despite the effective population size being explicitly accounted for in the substitution rates. Our results, along with the extensive literature on tree discordance due to incomplete lineage sorting [since it is closely linked to the effective population size; e.g., Maddison and Knowles (2006)], suggest that unaccounted-for variations in the effective population size may be the leading cause of errors in phylogenetic inference between closely related taxa.

The agnosticism of the substitution models with respect to the population size is also observed in studies that superimpose population genetic formulae on the substitution rates. While these studies have been able to quite successfully estimate mutation and selection bias [e.g., Lartillot (2013); Latrille and Lartillot (2022)], inferring variations in the population size has been more challenging. To circumvent this limitation, proxies for population size, such as life-history traits (Romiguier et al., 2014; Ellegren and Galtier, 2016) or polymorphic data are incorporated (Brevet and Lartillot, 2021), or further assumptions about the demographic history along the phylogeny are made [e.g., Latrille et al. (2021) assumes an autocorrelated geometric Brownian process to model variations in the effective population size]. These studies show that the patterns of nucleotide substitution are not *per se* sufficient to estimate variations in the effective population.

It is important to note that the population dynamics modeled in this study are still a far cry from reality. For example, phylogenetic incongruence due to ancient or recent gene flow might be another important source of error. Gene flow is likely to have a considerable impact on genetic distances, as it is pervasive between closely related taxa and during speciation (Pinho and Hey, 2010). Future work will include investigating this critical force. Nonetheless, we anticipate that our findings will aid in the development of mutation-selection phylogenetic models and inspire integrative approaches between phylogenetic and population-genetic time scales.

## Materials and Methods

### Models for the substitution process

We present and contrast two alternative substitution models. The mathematical results for these models are thoroughly described in the supplementary text S1–S3.

### Genomic data

We collected genomic data from six biological taxa: great apes, mice, flycatchers, fruit flies, Arabidopsis, and baker’s yeast, totaling 4385 individuals, 179 populations, and 17 different species. Table 1 summarizes the type and source of genomic data used in this study. The number of individuals and sites for each population is given in table S1, and the sampled site frequency spectra for each population are provided as supplementary material. We used the software package VCFtools (Danecek et al., 2011) to extract information about individual genotypes from the VCF and GVCF files. For each genotype, the four nucleotide bases were recoded posteriorly to *W* (weak base, *A* or *T*) and *S* (strong base, *G* or *C*), and the site-frequency spectra of each population were calculated by counting the absolute frequency of *S* alleles at various genomic positions.

### Bayesian inferences with population genomic data

The stationary distribution of the Moran model with boundary mutations and selection [equations S8 and S9 of the text S2] represents the distribution of allele frequencies in a population and therefore is the population site-frequency spectrum. To model the distribution of the allele content in *M* allelic counts sampled from a population of *N* individuals, we used the stationary quantity in conjunction with a hypergeometric distribution. This model was then used to infer the mutation rates, selection coefficient, and population size of a given biallelic system using a Bayesian framework. We set the Markov Chain Monte Carlo algorithm using a Metropolis-Hastings step based on normal proposals. We used a gamma prior for the selection coefficient and a discrete gamma distribution for the population size. The average mutation rate is assumed to be known. The Bayesian estimator of the Site-frequency Spectrum (BESS) was implemented in C++ and is deposited on GitHub at https://github.com/mrborges23/BESS. At the same location, we provide a small tutorial explaining how to perform inferences with BESS. BESS takes as input allelic counts in a sample of genomic sites and empirically obtained mutation rates. The program also includes an estimator for the neutral Moran model with biased mutations [based on equation (S16)] that is several orders of magnitude faster than the one with selection.

For our genomic data, we have used the sampled counts for the weak-strong alleles, along with the mutation rates provided in table 1. To estimate the population parameters, we used BESS in two steps: first, we used the neutral model to estimate the population size; second, we used the model with selection, employing the posterior distribution of the population size as priors. The parameter estimates for the 179 populations are provided as supplementary material.

### Sensitivity analyses

We conducted a sensitivity analysis to quantify the effect of different population forces on the measures of rate, time, and allele composition under the two models. In particular, we varied each population parameter *θ* by 10% and calculated the relative difference between the perturbed and unperturbed estimate *g* of the divergence rate, substitution time, or excess of fixed sites. The relative difference was calculated via the formula

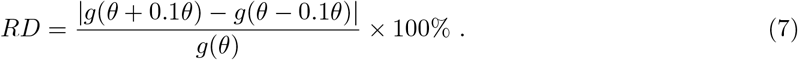

### Statistical analyses

The statistical analyses were performed using R version 4.1.2 (R Core Team, 2021). Because the number of populations varies across taxa and we wanted to avoid biasing our correlations by well-sampled taxa (e.g., yeasts are represented by 82 populations), we computed the correlation coefficients *ρ*_*B*_ between any two variables using the bootstrap technique (Li and Zharkikh, 1994). The correlation coefficient distribution was obtained by randomly sampling one variable value per taxon at each draw. We used 1000 bootstraps; the resulting distributions are in figures S6–S8. We then calculated the quantity

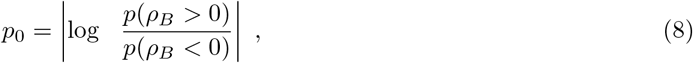

which is zero when the distribution is symmetric around zero. We set *p*_0_ *>* 2.944 as the significance threshold, which expresses the situation where *ρ*_*B*_ is greater or less than 0 with a probability of 0.95. Despite correcting for sampling bias, we have not performed a correction for the phylogenetic effects because we intended also to include the impact of the evolutionary relationships among the taxa. All correlations were performed with log-transformed variables.

## Acknowledgments

We thank all members of the Institute of Population Genetics for their feedback and support. This research was funded by the Austrian Science Fund (FWF) [P34524-B] to RB.

## Authors’ contributions

RB and CK conceived the idea. RB implemented the models, analyzed the data, and led the writing of the manuscript. IK, JB, MC, CFM, and CK collected data and contributed to the data analysis. All authors contributed critically to the drafts and gave their final approval for publication.

## Data availability

The data used and produced in this manuscript are available at https://vetcloud.vetmeduni.ac.at/nextcloud/s/JNMRXkKAn65ScHy and will be moved to Zenodo at a later stage.

## References

Bergman, J. and Schierup, M. H. (2021). Population dynamics of gc-changing mutations in humans and great apes. Genetics, 218.

Bolívar, P., Guéguen, L., Duret, L., Ellegren, H., and Mugal, C. F. (2019). Gc-biased gene conversion conceals the prediction of the nearly neutral theory in avian genomes. Genome Biology, 20:5.

Bolívar, P., Mugal, C. F., Nater, A., and Ellegren, H. (2016). Recombination rate variation modulates gene sequence evolution mainly via gc-biased gene conversion, not hill–robertson interference, in an avian system. Molecular Biology and Evolution, 33:216–227.

Borges, R., Boussau, B., Szöllősi, G. J., and Kosiol, C. (2022). Nucleotide usage biases distort inferences of the species tree. Genome Biology and Evolution, 14.

Borges, R. and Kosiol, C. (2020). Consistency and identifiability of the polymorphism-aware phylogenetic models. Journal of Theoretical Biology, 486:110074.

Borges, R., Szöllősi, G. J., and Kosiol, C. (2019a). Quantifying GC-Biased Gene Conversion in Great Ape Genomes Using Polymorphism-Aware Models. Genetics, 212(4):1321–1336.

Borges, R., Szöllősi, G., and Kosiol, C. (2019b). Quantifying gc-biased gene conversion in great ape genomes using polymorphism-aware models. Genetics, page genetics.302074.2019.

Brandvain, Y. and Wright, S. I. (2016). The limits of natural selection in a nonequilibrium world. Trends in Genetics, 32:201–210.

Brevet, M. and Lartillot, N. (2021). Reconstructing the history of variation in effective population size along phylogenies. Genome Biology and Evolution, 13.

Burri, R., Nater, A., Kawakami, T., Mugal, C. F., Olason, P. I., Smeds, L., Suh, A., Dutoit, L., Bureš, S., Garamszegi, L. Z., Hogner, S., Moreno, J., Qvarnström, A., Ružić, M., Sæther, S.-A., Sætre, G.-P., Török, J., and Ellegren, H. (2015). Linked selection and recombination rate variation drive the evolution of the genomic landscape of differentiation across the speciation continuum of ¡i¿ficedula¡/i¿ flycatchers. Genome Research, 25:1656–1665.

Chase, M. A. and Mugal, C. F. (2022). The role of recombination dynamics in shaping signatures of direct and indirect selection across the ficedula flycatcher genome. bioRxiv.

Cvijović, I., Good, B. H., and Desai, M. M. (2018). The effect of strong purifying selection on genetic diversity. Genetics, 209:1235–1278.

Danecek, P., Auton, A., Abecasis, G., Albers, C. A., Banks, E., DePristo, M. A., Handsaker, R. E., Lunter, G., Marth, G. T., Sherry, S. T., McVean, G., and Durbin, R. (2011). The variant call format and vcftools. Bioinformatics, 27:2156–2158.

De Maio, N., Schlötterer, C., and Kosiol, C. (2013). Linking great apes genome evolution across time scales using polymorphism-aware phylogenetic models. Molecular Biology and Evolution, 30(10):2249–2262.

De Maio, N., Schrempf, D., and Kosiol, C. (2015). PoMo: An allele frequency-based approach for species tree estimation. Systematic Biology, 64(6):1018–1031.

Duret, L. and Galtier, N. (2009). Biased gene conversion and the evolution of mammalian genomic landscapes. Annual Review of Genomics and Human Genetics, 10:285–311.

Ellegren, H. and Galtier, N. (2016). Determinants of genetic diversity. Nature Reviews Genetics, 17:422–433.

Fisher, R. (1930). The genetical theory of natural selection. Genetics, 154:272.

Flouri, T., Jiao, X., Rannala, B., and Yang, Z. (2020). A Bayesian Implementation of the Multispecies Coalescent Model with Introgression for Phylogenomic Analysis. Molecular Biology and Evolution, 37(4):1211–1223.

Galtier, N. (2021). Fine-scale quantification of gc-biased gene conversion intensity in mammals. Peer Community Journal, 1:e17.

Galtier, N., Roux, C., Rousselle, M., Romiguier, J., Figuet, E., Glémin, S., Bierne, N., and Duret, L. (2018). Codon usage bias in animals: Disentangling the effects of natural selection, effective population size, and gc-biased gene conversion. Molecular Biology and Evolution, 35:1092–1103.

Glémin, S., Arndt, P. F., Messer, P. W., Petrov, D., Galtier, N., and Duret, L. (2015). Quantification of GC-biased gene conversion in the human genome. Genome Research, 25(8):1215–1228.

Günther, T., Lampei, C., and Schmid, K. J. (2013). Mutational bias and gene conversion affect the intraspecific nitrogen stoichiometry of the arabidopsis thaliana transcriptome. Molecular Biology and Evolution, 30:561–568.

Halpern, A. L. and Bruno, W. J. (1998). Evolutionary distances for protein-coding sequences: modeling site-specific residue frequencies. Molecular Biology and Evolution, 15(7):910–917.

Harr, B., Karakoc, E., Neme, R., Teschke, M., Pfeifle, C., Željka Pezer, Babiker, H., Linnenbrink, M., Montero, I., Scavetta, R., Abai, M. R., Molins, M. P., Schlegel, M., Ulrich, R. G., Altmüller, J., Franitza, M., Büntge, A., Künzel, S., and Tautz, D. (2016). Genomic resources for wild populations of the house mouse, mus musculus and its close relative mus spretus. Scientific Data, 3:160075.

Hasegawa, M., Kishino, H., and Yano, T. (1985). Dating of the human-ape splitting by a molecular clock of mitochondrial {DNA}. Journal of Molecular Evolution, 22:160–174.

Hervas, S., Sanz, E., Casillas, S., Pool, J. E., and Barbadilla, A. (2017). Popfly: the drosophila population genomics browser. Bioinformatics, 33:2779–2780.

Hodgins-Davis, A., Rice, D. P., and Townsend, J. P. (2015). Gene expression evolves under a house-of-cards model of stabilizing selection. Molecular Biology and Evolution, 32:2130–2140.

Initiative, A. G. (2000). Analysis of the genome sequence of the flowering plant arabidopsis thaliana. Nature, 408:796–815.

Jukes, T. H. and Cantor, C. R. (1969). Evolution of protein molecules. In Munro, H., editor, Mammalian Protein Metabolism, pages 21–132. Elsevier, New York.

Keightley, P. D., Trivedi, U., Thomson, M., Oliver, F., Kumar, S., and Blaxter, M. L. (2009). Analysis of the genome sequences of three ¡i¿drosophila melanogaster¡/i¿ spontaneous mutation accumulation lines. Genome Research, 19:1195–1201.

Kingman, J. F. C. (1982). On the genealogy of large populations. Journal of Applied Probability, 19(A):27–43.

Lartillot, N. (2013). Phylogenetic patterns of gc-biased gene conversion in placental mammals and the evolutionary dynamics of recombination landscapes. Molecular Biology and Evolution, 30:489–502.

Latrille, T., Lanore, V., and Lartillot, N. (2021). Inferring long-term effective population size with mutation–selection models. Molecular Biology and Evolution, 38:4573–4587.

Latrille, T. and Lartillot, N. (2022). An improved codon modeling approach for accurate estimation of the mutation bias. Molecular Biology and Evolution, 39.

Latrille, T., Rodrigue, N., and Lartillot, N. (2022). Genes and sites under adaptation at the phylogenetic scale also exhibit adaptation at the population-genetic scale. bioRxiv.

Leaché, A. D. and Oaks, J. R. (2017). The Utility of Single Nucleotide Polymorphism (SNP) Data in Phylogenetics. Annual Review of Ecology, Evolution, and Systematics, 48(1):69–84.

Lesecque, Y., Mouchiroud, D., and Duret, L. (2013). Gc-biased gene conversion in yeast is specifically associated with crossovers: Molecular mechanisms and evolutionary significance. Molecular Biology and Evolution, 30:1409–1419.

Li, W.-H. and Zharkikh, A. (1994). What is the bootstrap technique? Systematic Biology, 43:424–430.

Lynch, M. (2010). Rate, molecular spectrum, and consequences of human mutation. Proceedings of the National Academy of Sciences, 107:961–968.

Lynch, M., Ackerman, M. S., Gout, J.-F., Long, H., Sung, W., Thomas, W. K., and Foster, P. L. (2016). Genetic drift, selection and the evolution of the mutation rate. Nature Reviews Genetics, 17(11):704–714.

Lynch, M., Sung, W., Morris, K., Coffey, N., Landry, C. R., Dopman, E. B., Dickinson, W. J., Okamoto, K., Kulkarni, S., Hartl, D. L., and Thomas, W. K. (2008). A genome-wide view of the spectrum of spontaneous mutations in yeast. Proceedings of the National Academy of Sciences, 105:9272–9277.

Maddison, W. P. and Knowles, L. L. (2006). Inferring phylogeny despite incomplete lineage sorting. Systematic Biology, 55(1):21–30.

Moran, P. A. P. (1958). Random processes in genetics. Mathematical Proceedings of the Cambridge Philosophical Society, 54(1):60.

Müller, R., Kaj, I., and Mugal, C. F. (2022). A nearly neutral model of molecular signatures of natural selection after change in population size. Genome Biology and Evolution, 14.

Nagylaki, T. (1983). Evolution of a Finite Population under Gene Conversion. Proceedings of the National Academy of Sciences of the United States of America, 80(20):6278–6281.

Ohta, T. (1973). Slightly deleterious mutant substitutions in evolution. Nature, 246:96–98.

Ossowski, S., Schneeberger, K., Lucas-Lledó, J. I., Warthmann, N., Clark, R. M., Shaw, R. G., Weigel, D., and Lynch, M. (2010). The rate and molecular spectrum of spontaneous mutations in ¡i¿arabidopsis thaliana¡/i¿. Science, 327:92–94.

Pessia, E., Popa, A., Mousset, S., Rezvoy, C., Duret, L., and Marais, G. A. B. (2012). Evidence for widespread gc-biased gene conversion in eukaryotes. Genome Biology and Evolution, 4:675–682.

Peter, J., Chiara, M. D., Friedrich, A., Yue, J.-X., Pflieger, D., Bergström, A., Sigwalt, A., Barre, B., Freel, K., Llored, A., Cruaud, C., Labadie, K., Aury, J.-M., Istace, B., Lebrigand, K., Barbry, P., Engelen, S., Lemainque, A., Wincker, P., Liti, G., and Schacherer, J. (2018). Genome evolution across 1,011 saccharomyces cerevisiae isolates. Nature, 556:339–344.

Pinho, C. and Hey, J. (2010). Divergence with Gene Flow: Models and Data. Annual Review of Ecology, Evolution, and Systematics, 41(1):215–230.

R Core Team (2021). R: A Language and Environment for Statistical Computing. R Foundation for Statistical Computing, Vienna, Austria.

Rannala, B. and Yang, Z. (2003). Bayes estimation of species divergence times and ancestral population sizes using dna sequences from multiple loci. Genetics, 164:1645–1656.

Robinson, D., Jones, D., Kishino, H., Goldman, N., and Thorne, J. (2003). Protein Evolution with Dependence Among Codons Due to Tertiary Structure. Molecular Biology and Evolution, 20(10):1692–1704.

Robinson, M. C., Stone, E. A., and Singh, N. D. (2014). Population genomic analysis reveals no evidence for gc-biased gene conversion in drosophila melanogaster. Molecular Biology and Evolution, 31:425–433.

Romiguier, J., Gayral, P., Ballenghien, M., Bernard, A., Cahais, V., Chenuil, A., Chiari, Y., Dernat, R., Duret, L., Faivre, N., Loire, E., Lourenco, J. M., Nabholz, B., Roux, C., Tsagkogeorga, G., Weber, A. A.-T., Weinert, L. A., Belkhir, K., Bierne, N., Glémin, S., and Galtier, N. (2014). Comparative population genomics in animals uncovers the determinants of genetic diversity. Nature, 515:261–263.

Schrempf, D., Minh, B. Q., De Maio, N., von Haeseler, A., and Kosiol, C. (2016). Reversible polymorphism-aware phylogenetic models and their application to tree inference. Journal of Theoretical Biology, 407:362–370.

Smeds, L., Qvarnström, A., and Ellegren, H. (2016). Direct estimate of the rate of germline mutation in a bird. Genome Research, 26:1211–1218.

Tavaré, S. (1986). Some Probabilistic and Statistical Problems in the Analysis of DNA Sequences. Lectures on Mathematics in the Life Sciences, 17:57–86.

Uchimura, A., Higuchi, M., Minakuchi, Y., Ohno, M., Toyoda, A., Fujiyama, A., Miura, I., Wakana, S., Nishino, J., and Yagi, T. (2015). Germline mutation rates and the long-term phenotypic effects of mutation accumulation in wild-type laboratory mice and mutator mice. Genome Research, 25:1125–1134.

Venn, O., Turner, I., Mathieson, I., de Groot, N., Bontrop, R., and McVean, G. (2014). Strong male bias drives germline mutation in chimpanzees. Science, 344:1272–1275.

Wright, S. (1931). Evolution in mendelian populations. Genetics, 16:97–159.

