## supplemental text, tables and figures for "Traditional phylogenetic models fail to account for variations in the effective population size"

### Supplementary Files

#### Supplementary Text. Mathematical results

List of parameters:

- $\lambda_{AB}$: the $A$ to $B$ substitution rate, here constrained between $[0, 1]$ and defined as the probability of a substitution from allele $A$ to $B$ per time unit
- $\lambda_{BA}$: the $B$ to $A$ substitution rate, here constrained between $[0, 1]$ and defined as the probability of a substitution from allele $B$ to $A$ per time unit
- $\mu_{AB}$: the $A$ to $B$ mutation rate, here constrained between $[0, 1]$ and defined as the probability of a mutation from allele $A$ to $B$ per Moran event
- $\mu_{BA}$: the $B$ to $A$ mutation rate, here constrained between $[0, 1]$ and defined as the probability of a mutation from allele $B$ to $A$ per Moran event
- $\phi_A$: the fitness coefficient of allele $A$
- $\phi_B$: the fitness coefficient of allele $B$
- $N$: population size
- $p_{ij}$: the transition probability from state $i$ to $j$
- $s_i$: the stationary probability of state $i$
- $t_{ij}$: the time to reach state $j$ from $i$ in Moran events
- $r$: the expected number of events in stationarity

#### Text S1. A Markov model for the substitution process

Consider a discrete-time Markov process of states $A$ and $B$. $p_{AB} = \lambda_{AB}$ and $p_{BA} = \lambda_{BA}$ are the transition probabilities of this chain and constitute the substitution probabilities between variants $A$ and $B$. Treatment of the general two-state discrete-time Markov chain can be easily found in several undergraduate texts in probability. Here, we revisited some of its mathematical results that are useful to characterize the time and pace of the substitution process.

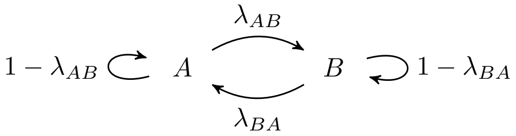


Figure S1: **The substitution process.** A model for the substitution process between any two variants represented by the states $A$ and $B$. $\lambda_{AB}$ and $\lambda_{BA}$ are the probabilities that a substitution between variants $A$ to $B$ occurs at each time step.

**The expected time of a substitution.** The expected time of a substitution is the time of the first arrival in $B$ from $A$ (or the other way around). Let $t_{AB}$ be the expected time to reach state $B$ if we started from state $A$. We know that

$$t_{AB} = 1 + (1 - \lambda_{AB})t_{AB} + \lambda_{AB}t_{BB} \quad . \tag{S1}$$

By definition $t_{BB} = 0$ and thus we obtain that $t_{AB} = \frac{1}{\lambda_{AB}}$. Similarly, $t_{BA} = \frac{1}{\lambda_{BA}}$.

**The stationary distribution.** Because this Markov chain is reversible, we can write that

$$s_A p_{AB} = s_B p_{BA} \quad , \tag{S2}$$

where $s_A$ are $s_B$ are elements of the stationary distribution. Since we know that $s_A + s_B = 1$, we obtain that $s_A = \frac{\lambda_{BA}}{\lambda_{AB}+\lambda_{BA}}$ and $s_B = \frac{\lambda_{AB}}{\lambda_{AB}+\lambda_{BA}}$.

**The expected divergence.** The expected divergence is the expected number of substitutions per unit of time. This value depends on the frequency of $A$ and $B$, which in our Markov process might vary over time. Here, we will calculate the expected divergence at steady state, for which we know the frequencies of $A$ and $B$. The expected divergence is defined by

$$d = s_A\lambda_{AB} + s_B\lambda_{BA} \quad , \tag{S3}$$

from which we can conclude that $d = 2\frac{\lambda_{AB}\lambda_{BA}}{\lambda_{AB}+\lambda_{BA}}$.

**Similarities with the time-continuous substitution model.** In phylogenetics, one usually models sequence evolution using time-continuous Markov chains. Here, we used a time-discrete one in order to make it comparable with the Moran model. If we redefine the substitution rates $\lambda_{AB}$ and $\lambda_{BA}$ so they represent the instantaneous substitutions rates, one can define this time-continuous process by the rate matrix

$$Q = \{q_{ij}\} = \begin{bmatrix} -\lambda_{AB} & \lambda_{AB} \\ \lambda_{BA} & -\lambda_{AB} \end{bmatrix} \quad . \tag{S4}$$

By solving the system of equations $sQ = 0$, one can verify that its stationary distribution is equal to the discrete-time process. In addition, the expected divergence, defined for the time-continuous process as

$$d = -s_A q_{AA} - s_B q_{BB} = 2\frac{\lambda_{AB}\lambda_{BA}}{\lambda_{AB} + \lambda_{BA}} \quad , \tag{S5}$$

also matches the expected divergence of the discrete model. The expected time to substitution is also the same between models, though, in the continuous-time model, the substitution time follows an exponential distribution instead of a geometric distribution. Therefore, the results of the discrete-time process are directly comparable with the time-continuous one.

#### Text S2. The Moran model with reversible mutation and selection

Consider a general Markov process on states $0, 1, 2, \ldots, N$, where the states represent the frequency of the variant $B$ in a population of $N$ individuals. When the population is at state 0 all the individuals have the variant $A$, whereas all the individuals have variant $B$ when the population is at state $N$. These states are known as fixed, boundary or monomorphic. The states between 1 and $N - 1$ are polymorphic, meaning that both alleles coexist in the population: i.e., some individuals have the variant $A$ while the remaining ones have the variant $B$.

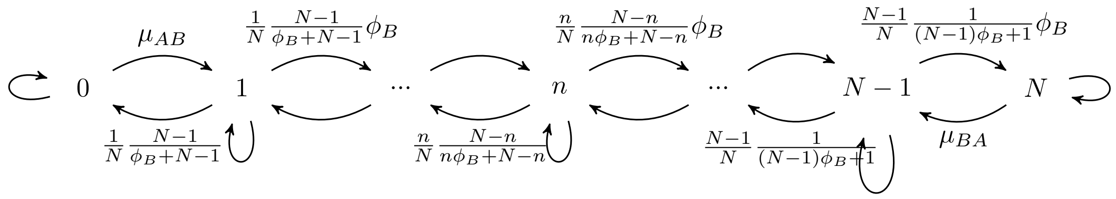


Figure S2: **A Moran model with boundary mutations and selection.** A model for the substitution process between two variants, $A$ and $B$. The states represent the frequency of the $B$ allele. $\{0\}$ and $\{N\}$ represent fixed states, and all the remaining ones are polymorphic. $\mu_{AB}$ and $\mu_{BA}$ represent the probabilities of a mutation from variant $A$ to $B$ and $B$ to $A$, respectively. In the Moran model, $\frac{n}{N}\frac{N-n}{N}$ represents the probability of picking an allele to die and an allele to reproduce. In this figure, this ratio is adapted to account for selection, where $\phi_B$ is the fitness coefficient on $B$. For the sake of clarity, the probabilities of remaining in the same state were not represented.

The probability of a state change is determined by a Moran model (Moran, 1958) with boundary mutations (mutations can only occur when one of the variants is fixed), genetic drift, and selection.

- $\mu_{AB}$ is the probability that a mutation from variant $A$ to $B$ occurs and vice versa to $\mu_{BA}$. The mutation probabilities are not necessarily the same, so the model accounts for possible mutation bias. Furthermore, we assume reversible mutations, so this model does not make the infinite site assumption. This assumption would limit the comparison with the traditional substitution models, which implicitly assume the reversibility of mutations.

- Selection is modeled via the fitness coefficient $\phi_B$ of variant $B$ that provides a reproductive advantage or disadvantage ($\phi_B > 1$ or $\phi_B < 1$, respectively) to the individuals possessing it. Selection acts to promote the fixation of $B$ variants.

- Genetic drift follows the Moran model, where at each generation, one random individual is chosen to die, and one random individual is chosen to reproduce. The strength of genetic drift is thus determined by the population size, which in our model is $N$.

Overall, the probability matrix of this Markov process is

$$P = \{p_{ij}\} = \begin{cases} \mu_{AB} & \text{if } i = 0 \text{ and } j = 1 \\ \mu_{BA} & \text{if } i = N \text{ and } j = N-1 \\ \frac{n(N-n)}{N(n\phi_B+N-n)}\phi_B & \text{if } i = n \text{ and } j = n+1 \\ \frac{n(N-n)}{N(n\phi_B+N-n)} & \text{if } i = n \text{ and } j = n-1 \\ 0 & \text{if } |i-j| > 1 \end{cases} , \tag{S6}$$

where the diagonal elements were defined such that $p_{ii} = 1 - \sum_{j \neq i} p_{ij}$. Some of the mathematical results that follow can be found in Borges et al. (2022, 2019b); they are here adapted for the biallelic case.

**The stationary distribution.** Because this Markov chain is reversible, we can use the detailed balance equations to obtain the stationary distribution: i.e., $s_i p_{ij} = s_j p_{ji}$. These equations allow us to obtain the frequencies of every state in terms of that of state 0. For $0 < n < N$, we have the recursion

$$s_n = s_0 \frac{N}{n} \frac{n\phi_B + N - n}{N - n} \mu_{AB} \phi_B^{n} - 1 \ , \tag{S7}$$

while for $s_N = s_0 \frac{\mu_{AB}}{\mu_{BA}} \phi_B^{N-1}$. By applying the constraint that the stationary frequencies have to sum up to one, we can derive expressions for the stationary frequencies for the boundary states

$$s_0 = \mu_{BA}\frac{1}{k} \quad \text{and} \quad s_N = \mu_{AB}\phi_B^{N-1}\frac{1}{k} \quad , \tag{S8}$$

and polymorphic states

$$s_n = \mu_{AB}\mu_{BA}\phi_B^{n-1}\frac{N}{n}\frac{n\phi_B + N - n}{N-n}\frac{1}{k} \quad , \tag{S9}$$

where $k$ is the normalization constant

$$k = \mu_{AB}\phi_B^{N-1} + \mu_{BA} + \mu_{AB}\mu_{BA}\sum_{n=1}^{N-1}\phi_B^{n-1}\frac{N}{n}\frac{n\phi_B + N - n}{N-n} \quad . \tag{S10}$$

**The expected time of a substitution.** The expected time of a substitution is the time of the first arrival in $N$ from 0 (or the other way around). Let $t_{0N}$ be the expected time to reach state 0 if we started from state $N$. We use the definition of first passage time in a Markov of chain, which uses the idea to condition on where the chain goes after one transition: i.e.,

$$t_{0N} = 1 + \sum_{k=0}^{N} p_{0k}t_{kN} \quad . \tag{S11}$$

From this, we obtained the recursion

$$t_{nN} = t_{n+1N} + \frac{1}{\mu_{AB}\phi_B^n} + N\sum_{k=1}^{n}\frac{k\phi_B + N - k}{k(N-k)\phi_B^{n-k+1}} \quad . \tag{S12}$$

From this recursion, we can determine $t_{N-1\ N}$ as by definition $t_{NN} = 0$. As one element is now defined, we now use the recursion backwards and derive

$$t_{0N} = \frac{1}{\mu_{AB}}\sum_{n=0}^{N-1}\frac{1}{\phi_B^n} + N\sum_{k=1}^{N-1}\sum_{n=k}^{N-k}\frac{n\phi_B + N - n}{n(N-n)\phi_B} \quad . \tag{S13}$$

For the neutral case ($\phi_B = 1$), this expression can be significantly simplified to obtain

$$t_{0N} = \frac{N}{\mu_{AB}} + N^2 H_{N-1} \quad , \tag{S14}$$

where $H_{N-1} = \sum_{n=1}^{N-1}\frac{1}{n}$ is an harmonic number.

**The expected divergence.** As for the substitution process, we will calculate the expected divergence at the steady state. The expected number of events is given by

$$s_0\mu_{AB} + s_N\mu_{BA} + \sum_{n=1}^{N-1} s_n\left[\frac{n(N-n)}{N[n\phi_B + N - n]} + \frac{n(N-n)}{N[n\phi_B + N - n]}\phi_B\right] \quad , \tag{S15}$$

which by substituting the stationary frequencies of each state can be simplified to

$$2\mu_{AB}\mu_{BA}\frac{1}{k}\sum_{n=0}^{N-1}\phi_B^n \quad . \tag{S16}$$

For the neutral case (i.e., $\phi_B = 1$), we can further simplify the initial expression to

$$2\mu_{AB}\mu_{BA}N\frac{1}{k} \quad . \tag{S17}$$

These quantities express the expected number of mutations and frequency shifts per Moran event. If we further divide them by $N$, we obtain the expected divergence per Moran generations. This is the quantity that we will mention in the main text.

#### Text S3. Sampling alleles from the population

Consider that we sample $M$ individuals from a population of $N$ individuals. Let $m$ be the number of $B$ variants counted among the $M$ individuals and $n$ the number of $B$ variants in the original population of size $N$. $m$ varies between 0 and $M$: if $m = 0$ all the sampled individuals have the variant $A$; if $m = M$ all the sampled individuals have the variant $B$; if $0 < m < N$, $m$ individuals have the variant $B$ and $M - m$ the variant $A$. For obvious reasons, the first two types of counts are called monomorphic, while the latter is called polymorphic.

If one observes a polymorphic count, then the original population from which the count was sampled must be polymorphic. The probability $c_m$ of observing such a count is the probability of the population being at the polymorphic state $n$ times the probability of sampling $m$ $B$ variants and $M - m$ $A$ variants from that population (provided that $n \geq m$ and $N - n \geq M - m$). The probability of the population being at the polymorphic state $nB$, will vary through time. As we assumed for calculating the expected divergence, we will assume the population is close or at the stationary phase. The mentioned probability then becomes,

$$c_m = \sum_{n=m}^{N-M+m} s_n \left[ \prod_{i=0}^{m-1} \frac{n-i}{N-i} \right] \left[ \prod_{i=0}^{M-m-1} \frac{N-n-i}{N-m-i} \right] \binom{M}{m} \ . \tag{S18}$$

If one observes a monomorphic count of type $B$, then it is less obvious to determine whether the original population is fixed for $B$, or is in a polymorphic state where $B$ is frequent enough to generate monomorphic counts. The probability $c_M$ of observing a monomorphic count of type $B$ is the sum of the probability of extracting $M$ $B$ alleles from a population with $N$ $B$ alleles or any of the polymorphic states where $B$ is present and its frequency $n \geq M$.

$$c_M = s_N + \sum_{n=M}^{N-1} s_n \prod_{i=0}^{M-1} \frac{n-i}{N-i} \tag{S19}$$

The last term of the equation expresses the degree to which the monomorphic counts are overestimated. As expected, this term converges to zero as $M$ approaches $N$.

**Establishing a relationship between the substitution and the Moran model**: The use of a single sequence per species in phylogenetic analyses, can be seen, on a population genetics framework, as sampling a single individual from the population. This result has two possible outcomes: an individual of type $A$ or $B$. The probabilities of observing such individuals can be easily calculated by adapting the equation (S19) for the situation when $M = 1$:

$$c_1 = \sum_{n=1}^{N} s_n \frac{n}{N} = \frac{1}{k}\mu_{AB}\phi_B^{N-1} + \frac{1}{k}\mu_{AB}\mu_{BA}\sum_{n=1}^{N-1}\phi_B^{n-1}\frac{n\phi_B + N - n}{N-n} \tag{S20}$$

$$= \frac{1}{k}\mu_{AB}\phi_B^{N-1} + \frac{1}{k}\mu_{AB}\mu_{BA}\left[1 - \phi_B^{N-1} + N\sum_{n=1}^{N-1}\frac{\phi_B^{N-n}}{n}\right] \quad \text{and}$$

$$c_0 = \sum_{n=0}^{N-1} s_n \frac{N-n}{N} = \frac{1}{k}\mu_{BA} + \frac{1}{k}\mu_{AB}\mu_{BA}\sum_{n=1}^{N-1}\phi_B^{n-1}\frac{n\phi_B + N - n}{n} \tag{S21}$$

$$= \frac{1}{k}\mu_{BA} + \frac{1}{k}\mu_{AB}\mu_{BA}\left[\phi_B^{N-1} - 1 + N\sum_{n=1}^{N-1}\frac{\phi_B^{n-1}}{n}\right] .$$

In the main text, $c_0 = p_A$ and $c_1 = p_B$.

To establish a relation between the substitution rates and these probabilities, we assume that these probabilities should match the expected frequencies of the states $W$ and $S$ in the substitution model. In stationarity, these frequencies are known, which allow us to write:

$$p_A = s_A = \frac{\lambda_{BA}}{\lambda_{AB} + \lambda_{BA}} \quad \text{and} \quad p_B = s_B = \frac{\lambda_{AB}}{\lambda_{AB} + \lambda_{BA}} . \tag{S22}$$

However, we cannot solve this equations to $\lambda_{BA}$ and $\lambda_{BA}$, because we have no information about the value of $\lambda_{AB} + \lambda_{BA}$. The unidentifiability of the substitution rates from the frequencies of $S$ and $W$ is the reason why genetic distances under the phylogenetic substitution models are usually normalized so that one substitution per unit of time is expected. We thus define instead the relations

$$\lambda_{AB} \propto \mu_{AB}\phi_B^{N-1} + \mu_{AB}\mu_{BA}\left[1 - \phi_B^{N-1} + N\sum_{n=1}^{N-1}\frac{\phi_B^{N-n}}{n}\right] \quad \text{and} \tag{S23}$$

$$\lambda_{BA} \propto \mu_{BA} + \mu_{AB}\mu_{BA}\left[\phi_B^{N-1} - 1 + N\sum_{n=1}^{N-1}\frac{\phi_B^{n-1}}{n}\right] , \tag{S24}$$

which cannot be interpreted in terms of their absolute values, but still allow testing the relative impact of the population parameters on the substitution rates.

### Supplementary Tables

Table S1: **Description of the population genomic data.** Legend: number of individuals (or the number of haploid individuals for the diploid organisms; $NI$), number of sites ($NS$), proportion of monomorphic sites of type week ($PW$), proportion of monomorphic sites of type strong ($PS$) and proportion of polymorphic sites ($PP$).

| **Taxa** | **Population** | $NI$ | $NS$ | $PW$ | $PS$ | $PP = 1 - PW - PS$ |
|---|---|---|---|---|---|---|
| great_apes | GBB | 14 | 9999889 | 0.742 | 0.257 | 0.00051 |
| great_apes | GBG | 16 | 9999621 | 0.756 | 0.244 | 0.00058 |
| great_apes | GGG | 46 | 9996329 | 0.756 | 0.242 | 0.00172 |
| great_apes | PAB | 22 | 9996105 | 0.751 | 0.247 | 0.00201 |
| great_apes | PPA | 26 | 9998870 | 0.753 | 0.246 | 0.00108 |
| great_apes | PPY | 30 | 9996700 | 0.755 | 0.243 | 0.00146 |
| great_apes | PTE | 20 | 9998847 | 0.74 | 0.258 | 0.00144 |
| great_apes | PTS | 38 | 9997618 | 0.748 | 0.25 | 0.00187 |
| great_apes | PTT | 36 | 9997671 | 0.748 | 0.249 | 0.00285 |
| great_apes | PTV | 22 | 9999059 | 0.746 | 0.253 | 0.00084 |
| great_apes | YRI | 44 | 9999999 | 0.738 | 0.26 | 0.00135 |
| mice | ms_spre | 16 | 26315005 | 0.576 | 0.416 | 0.00688 |
| mice | mmm_kaz | 16 | 26134928 | 0.578 | 0.417 | 0.00469 |
| mice | mmm_cze | 16 | 26116234 | 0.578 | 0.417 | 0.00465 |
| mice | mmm_afg | 12 | 26124859 | 0.577 | 0.416 | 0.00567 |
| mice | mmd_ira | 16 | 26156328 | 0.575 | 0.415 | 0.00898 |
| mice | mmd_hel | 6 | 25954326 | 0.580 | 0.418 | 0.00122 |
| mice | mmd_ger | 16 | 26028499 | 0.578 | 0.416 | 0.00473 |
| mice | mmd_fra | 16 | 26044333 | 0.578 | 0.416 | 0.00505 |
| mice | mmc_cast | 20 | 26450657 | 0.569 | 0.410 | 0.01937 |
| flycatchers | coll | 190 | 1824118 | 0.543 | 0.447 | 0.00824 |
| flycatchers | par | 30 | 1804535 | 0.546 | 0.444 | 0.00848 |
| flycatchers | pied | 22 | 1669283 | 0.565 | 0.429 | 0.00468 |
| flycatchers | taig | 130 | 1738330 | 0.552 | 0.432 | 0.01518 |
| flycatchers | semi | 40 | 1534463 | 0.581 | 0.412 | 0.00575 |
| flycatchers | spec | 40 | 1794735 | 0.550 | 0.444 | 0.00521 |
| drosophila | AUS | 13 | 868469 | 0.409 | 0.578 | 0.01309 |
| drosophila | CHB | 11 | 706971 | 0.425 | 0.565 | 0.00935 |
| drosophila | CO | 10 | 1474317 | 0.428 | 0.555 | 0.01663 |
| drosophila | EA | 9 | 649941 | 0.426 | 0.561 | 0.01344 |
| drosophila | EB | 5 | 2759137 | 0.432 | 0.558 | 0.01002 |
| drosophila | ED | 6 | 1583955 | 0.42 | 0.57 | 0.01026 |
| drosophila | EF | 40 | 183622 | 0.424 | 0.551 | 0.02499 |
| drosophila | EG | 11 | 419943 | 0.432 | 0.557 | 0.01053 |
| drosophila | ER | 5 | 2972444 | 0.43 | 0.56 | 0.00935 |
| drosophila | EZ | 4 | 1514712 | 0.432 | 0.558 | 0.00955 |
| drosophila | FR | 69 | 244482 | 0.442 | 0.54 | 0.01815 |
| drosophila | GA | 9 | 1506833 | 0.428 | 0.555 | 0.01644 |
| drosophila | GU | 5 | 1488725 | 0.431 | 0.557 | 0.01208 |
| drosophila | KN | 5 | 1505651 | 0.431 | 0.557 | 0.0118 |
| drosophila | KR | 4 | 1502541 | 0.432 | 0.558 | 0.00951 |
| drosophila | MW | 6 | 1397937 | 0.428 | 0.558 | 0.01371 |
| drosophila | NG | 6 | 2959404 | 0.429 | 0.559 | 0.01248 |
| drosophila | NTH | 17 | 558990 | 0.403 | 0.586 | 0.01148 |
| drosophila | RAL | 177 | 169141 | 0.426 | 0.545 | 0.02921 |
| drosophila | RG | 27 | 2877618 | 0.421 | 0.554 | 0.02445 |
| drosophila | SB | 5 | 2959107 | 0.429 | 0.558 | 0.01259 |
| drosophila | SD | 20 | 332360 | 0.422 | 0.56 | 0.01819 |
| drosophila | SF | 4 | 1588382 | 0.427 | 0.561 | 0.01164 |
| drosophila | SP | 19 | 489657 | 0.42 | 0.554 | 0.02643 |

Table S1: (continued from previous page)

| **Taxa** | **Population** | $NI$ | $NS$ | $PW$ | $PS$ | $PP = 1 - PW - PS$ |
|---|---|---|---|---|---|---|
| drosophila | UG | 4 | 1502343 | 0.432 | 0.558 | 0.01065 |
| drosophila | UK | 4 | 1549094 | 0.429 | 0.56 | 0.01121 |
| drosophila | USI | 13 | 620431 | 0.428 | 0.559 | 0.01281 |
| drosophila | USW | 22 | 337640 | 0.428 | 0.557 | 0.01532 |
| drosophila | ZI | 197 | 2386074 | 0.395 | 0.543 | 0.0612 |
| drosophila | ZS | 4 | 2112544 | 0.43 | 0.558 | 0.0113 |
| arabidopsis | ARM | 7 | 58265314 | 0.609 | 0.389 | 0.00159 |
| arabidopsis | AUT | 13 | 57721467 | 0.609 | 0.387 | 0.0048 |
| arabidopsis | AZE | 16 | 56533275 | 0.612 | 0.386 | 0.0019 |
| arabidopsis | BEL | 2 | 58364468 | 0.609 | 0.389 | 0.00229 |
| arabidopsis | BUL | 28 | 56119489 | 0.613 | 0.384 | 0.00341 |
| arabidopsis | CAN | 2 | 58385798 | 0.609 | 0.389 | 0.00223 |
| arabidopsis | CRO | 2 | 58356005 | 0.609 | 0.389 | 0.0019 |
| arabidopsis | CZE | 40 | 55491763 | 0.616 | 0.383 | 0.00105 |
| arabidopsis | DEN | 2 | 58375613 | 0.609 | 0.389 | 0.00241 |
| arabidopsis | ESP | 180 | 55122169 | 0.616 | 0.382 | 0.00117 |
| arabidopsis | FIN | 3 | 58097410 | 0.609 | 0.389 | 0.00204 |
| arabidopsis | FRA | 45 | 55735992 | 0.614 | 0.383 | 0.00217 |
| arabidopsis | GEO | 14 | 57072833 | 0.611 | 0.386 | 0.00297 |
| arabidopsis | GER | 118 | 55201186 | 0.617 | 0.383 | 0.00086 |
| arabidopsis | GRC | 3 | 58149770 | 0.609 | 0.389 | 0.00248 |
| arabidopsis | ITA | 73 | 55022850 | 0.617 | 0.383 | 0.00028 |
| arabidopsis | JPN | 2 | 58456523 | 0.609 | 0.389 | 0.00228 |
| arabidopsis | KAZ | 2 | 58432667 | 0.609 | 0.39 | 0.00075 |
| arabidopsis | KGZ | 5 | 58084320 | 0.609 | 0.389 | 0.00202 |
| arabidopsis | LBN | 2 | 58475108 | 0.608 | 0.388 | 0.00371 |
| arabidopsis | LTU | 4 | 58196350 | 0.609 | 0.388 | 0.00338 |
| arabidopsis | MAR | 2 | 58148653 | 0.609 | 0.39 | 0.00117 |
| arabidopsis | NED | 11 | 57542730 | 0.609 | 0.386 | 0.00509 |
| arabidopsis | NOR | 2 | 58579263 | 0.608 | 0.389 | 0.00262 |
| arabidopsis | POL | 3 | 58308954 | 0.608 | 0.388 | 0.00344 |
| arabidopsis | POR | 10 | 56880644 | 0.612 | 0.385 | 0.00275 |
| arabidopsis | ROU | 9 | 56713249 | 0.612 | 0.386 | 0.00228 |
| arabidopsis | RUS | 60 | 55434113 | 0.615 | 0.383 | 0.00104 |
| arabidopsis | SRB | 9 | 57248280 | 0.61 | 0.386 | 0.00324 |
| arabidopsis | SUI | 5 | 58127996 | 0.608 | 0.388 | 0.00403 |
| arabidopsis | SVK | 7 | 58028603 | 0.609 | 0.387 | 0.00402 |
| arabidopsis | SWE | 243 | 55458445 | 0.615 | 0.383 | 0.00179 |
| arabidopsis | TJK | 3 | 58359577 | 0.609 | 0.389 | 0.00141 |
| arabidopsis | UK | 69 | 55763042 | 0.615 | 0.384 | 0.00161 |
| arabidopsis | UKR | 2 | 58078741 | 0.609 | 0.389 | 0.00248 |
| arabidopsis | USA | 123 | 56010508 | 0.614 | 0.384 | 0.00196 |
| arabidopsis | UZB | 4 | 57297073 | 0.612 | 0.388 | 0.00027 |
| yeast | USA | 170 | 1544489 | 0.408 | 0.409 | 0.18295 |
| yeast | Ukraine | 4 | 1544489 | 0.49 | 0.497 | 0.01353 |
| yeast | UK | 42 | 1544489 | 0.43 | 0.436 | 0.1341 |
| yeast | Japan | 118 | 1544489 | 0.433 | 0.436 | 0.13115 |
| yeast | Slovakia | 50 | 1544489 | 0.454 | 0.461 | 0.0848 |
| yeast | Denmark | 6 | 1544489 | 0.476 | 0.483 | 0.04084 |
| yeast | Spain | 158 | 1544489 | 0.42 | 0.426 | 0.15423 |
| yeast | France | 186 | 1544489 | 0.413 | 0.417 | 0.16972 |
| yeast | Czech_Republic | 2 | 1544489 | 0.486 | 0.494 | 0.01949 |
| yeast | Australia | 12 | 1544489 | 0.475 | 0.483 | 0.04208 |
| yeast | Ireland | 2 | 1544489 | 0.496 | 0.504 | 0.00016 |
| yeast | South_Africa | 24 | 1544489 | 0.454 | 0.464 | 0.08135 |
| yeast | Finland | 12 | 1544489 | 0.472 | 0.48 | 0.04806 |

Table S1: (continued from previous page)

| Taxa | Population | $NI$ | $NS$ | $PW$ | $PS$ | $PP = 1 - PW - PS$ |
|---|---|---|---|---|---|---|
| yeast | Italy | 210 | 1544489 | 0.421 | 0.429 | 0.14936 |
| yeast | Brazil | 86 | 1544489 | 0.443 | 0.449 | 0.10796 |
| yeast | China | 28 | 1544489 | 0.411 | 0.41 | 0.1785 |
| yeast | Philippines | 12 | 1544489 | 0.457 | 0.464 | 0.07901 |
| yeast | Ivory_Coast | 16 | 1544489 | 0.45 | 0.462 | 0.08783 |
| yeast | Indonesia | 16 | 1544489 | 0.463 | 0.469 | 0.06825 |
| yeast | Austria | 8 | 1544489 | 0.479 | 0.486 | 0.03481 |
| yeast | Russia | 24 | 1544489 | 0.458 | 0.465 | 0.07661 |
| yeast | Netherlands | 32 | 1544489 | 0.442 | 0.448 | 0.10953 |
| yeast | Nigeria | 18 | 1544489 | 0.464 | 0.468 | 0.06826 |
| yeast | Vietnam | 26 | 1544489 | 0.455 | 0.461 | 0.084 |
| yeast | French_Guiana | 66 | 1544489 | 0.458 | 0.463 | 0.07882 |
| yeast | Sweden | 10 | 1544489 | 0.472 | 0.48 | 0.04828 |
| yeast | Puerto_Rico | 2 | 1544489 | 0.495 | 0.502 | 0.00315 |
| yeast | Pakistan | 2 | 1544489 | 0.496 | 0.504 | 0.00016 |
| yeast | Portugal | 10 | 1544489 | 0.484 | 0.491 | 0.02426 |
| yeast | Germany | 16 | 1544489 | 0.463 | 0.47 | 0.06735 |
| yeast | Chile | 12 | 1544489 | 0.489 | 0.496 | 0.01482 |
| yeast | Costa_Rica | 2 | 1544489 | 0.496 | 0.504 | 0.00015 |
| yeast | Switzerland | 6 | 1544489 | 0.488 | 0.495 | 0.017 |
| yeast | Taiwan | 34 | 1544489 | 0.421 | 0.42 | 0.15836 |
| yeast | Romania | 4 | 1544489 | 0.496 | 0.503 | 0.00079 |
| yeast | Ghana | 14 | 1544489 | 0.448 | 0.463 | 0.08939 |
| yeast | Cameroon | 6 | 1544489 | 0.492 | 0.499 | 0.0089 |
| yeast | West_Africa | 34 | 1544489 | 0.464 | 0.472 | 0.06433 |
| yeast | Malaysia | 14 | 1544489 | 0.475 | 0.482 | 0.04347 |
| yeast | Ecuador | 46 | 1544489 | 0.437 | 0.441 | 0.12235 |
| yeast | Burundi | 4 | 1544489 | 0.478 | 0.487 | 0.03463 |
| yeast | Navassa_Island | 2 | 1544489 | 0.485 | 0.493 | 0.02188 |
| yeast | Peru | 8 | 1544489 | 0.488 | 0.495 | 0.01634 |
| yeast | Korea | 4 | 1544489 | 0.485 | 0.493 | 0.02149 |
| yeast | Turkey | 4 | 1544489 | 0.478 | 0.486 | 0.03553 |
| yeast | South_Armenia | 2 | 1544489 | 0.496 | 0.503 | 0.00113 |
| yeast | Bulgaria | 4 | 1544489 | 0.479 | 0.487 | 0.0342 |
| yeast | Belgium | 18 | 1544489 | 0.457 | 0.463 | 0.07967 |
| yeast | Czechoslovakia | 2 | 1544489 | 0.496 | 0.503 | 0.00108 |
| yeast | Hungary | 10 | 1544489 | 0.471 | 0.478 | 0.05085 |
| yeast | Pennsylvanian | 2 | 1544489 | 0.495 | 0.505 | 0.00017 |
| yeast | Djibouti | 16 | 1544489 | 0.477 | 0.485 | 0.03835 |
| yeast | Mexico | 18 | 1544489 | 0.473 | 0.48 | 0.04714 |
| yeast | Israel | 30 | 1544489 | 0.463 | 0.47 | 0.06634 |
| yeast | Equador | 2 | 1544489 | 0.486 | 0.494 | 0.02009 |
| yeast | Madagascar | 4 | 1544489 | 0.469 | 0.483 | 0.04789 |
| yeast | Chad | 10 | 1544489 | 0.482 | 0.495 | 0.02293 |
| yeast | Slovenia | 36 | 1544489 | 0.452 | 0.459 | 0.08893 |
| yeast | Serbia | 2 | 1544489 | 0.49 | 0.5 | 0.01008 |
| yeast | Montenegro | 2 | 1544489 | 0.496 | 0.503 | 0.00019 |
| yeast | Laos | 10 | 1544489 | 0.474 | 0.482 | 0.04449 |
| yeast | Croatia | 4 | 1544489 | 0.494 | 0.501 | 0.00581 |
| yeast | Greece | 6 | 1544489 | 0.491 | 0.498 | 0.01071 |
| yeast | Norway | 2 | 1544489 | 0.484 | 0.492 | 0.02337 |
| yeast | Ethiopia | 8 | 1544489 | 0.481 | 0.489 | 0.03027 |
| yeast | Malta | 2 | 1544489 | 0.496 | 0.503 | 0.00015 |
| yeast | Algeria | 2 | 1544489 | 0.494 | 0.503 | 0.00296 |
| yeast | Uruguay | 2 | 1544489 | 0.496 | 0.503 | 0.00014 |
| yeast | Argentina | 8 | 1544489 | 0.487 | 0.495 | 0.01775 |

Table S1: (continued from previous page)

| **Taxa** | **Population** | $NI$ | $NS$ | $PW$ | $PS$ | $PP = 1 - PW - PS$ |
|---|---|---|---|---|---|---|
| yeast | Canada | 2 | 1544489 | 0.495 | 0.504 | 2e-04 |
| yeast | Georgia | 52 | 1544489 | 0.481 | 0.489 | 0.02989 |
| yeast | Yugoslavia | 2 | 1544489 | 0.496 | 0.504 | 0.00017 |
| yeast | Sri_Lanka | 6 | 1544489 | 0.485 | 0.493 | 0.0225 |
| yeast | North_Korea | 4 | 1544489 | 0.485 | 0.494 | 0.02113 |
| yeast | Tailand | 4 | 1544489 | 0.494 | 0.503 | 0.0031 |
| yeast | Lebanon | 18 | 1544489 | 0.483 | 0.491 | 0.02621 |
| yeast | Northern_Europe | 4 | 1544489 | 0.474 | 0.482 | 0.04419 |
| yeast | Burkina_Faso | 8 | 1544489 | 0.483 | 0.494 | 0.02314 |
| yeast | Bahamas | 4 | 1544489 | 0.495 | 0.504 | 0.00133 |
| yeast | Hawaii | 4 | 1544489 | 0.495 | 0.503 | 0.00121 |
| yeast | Trinidad | 2 | 1544489 | 0.496 | 0.504 | 0.00019 |
| yeast | Jamaica | 2 | 1544489 | 0.496 | 0.504 | 0.00015 |
| yeast | Thailand | 4 | 1544489 | 0.495 | 0.503 | 0.00257 |
| yeast | Africa | 2 | 1544489 | 0.496 | 0.504 | 0.00016 |
| yeast | Phillipines | 2 | 1544489 | 0.496 | 0.504 | 0.00013 |
| yeast | India | 2 | 1544489 | 0.495 | 0.505 | 1e-04 |

Table S2: Per taxa correlation tests between the excess of fixed sites ($f$) and the proportion of polymorphic sites ($P$) and the sample size (or number of haploid individuals; $M$). The values in each cell represent the estimated Pearson's correlation coefficient $\rho$ and inside parenthesis the $p$-value of the test.

| **Taxa** | $\log(P)$ **vs.** $\log(f)$ | $\log(M)$ **vs.** $\log(f)$ |
|---|---|---|
| Great apes | 0.976 (<0.001) | 0.563 (0.071) |
| Mice | 0.995 (<0.001) | 0.808 (0.008) |
| Flycatchers | 0.540 (0.268) | 0.094 (0.860) |
| Fruit flies | −0.024 (0.900) | −0.416 (0.022) |
| Arabidopsis | 0.772 (<0.001) | −0.811 (<0.001) |
| Baker's yeast | 0.997 (<0.001) | 0.717 (<0.001) |

### Supplementary Figures

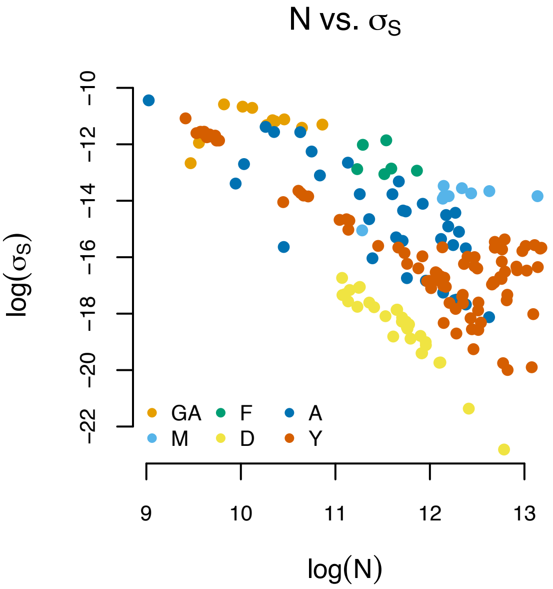


Figure S3: Correlation between the effective population size and the selection coefficient The correlation between the variables is strong and statistically significant: $\rho = -0.70$ and $p$–value<0.001.

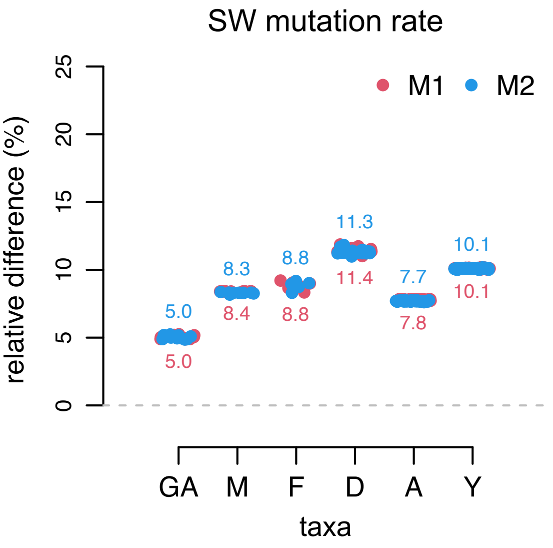


Figure S4: Sensitivity analyses for the SW mutation rate on the divergence rate. The relative difference expresses the effect of changing each population parameter by ±10% on the divergence rate. The numbers represent the average relative divergence across all populations. Legend: great apes (GA), mice (M), flycatchers (F), fruit flies (D), arabidopsis (A), and baker's yeast (Y).

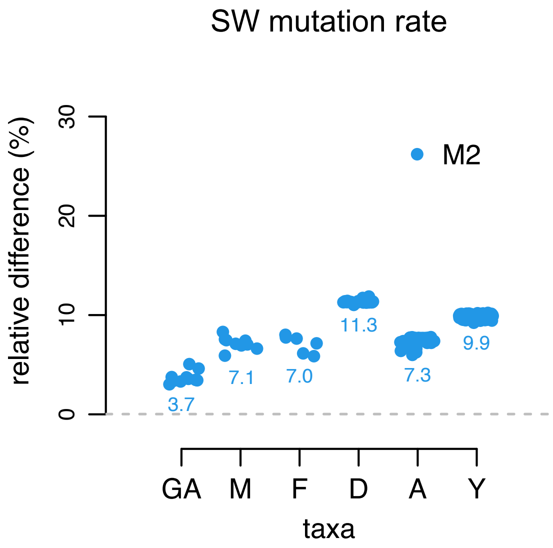


Figure S5: Sensitivity analyses for the SW mutation rate on the excess of fixed sites. The relative difference expresses the effect of changing each population parameter by ±10% on the excess of observed sites [equation (4)]. The numbers represent the average relative divergence across all populations. Legend: great apes (GA), mice (M), flycatchers (F), fruit flies (D), arabidopsis (A), and baker's yeast (Y).

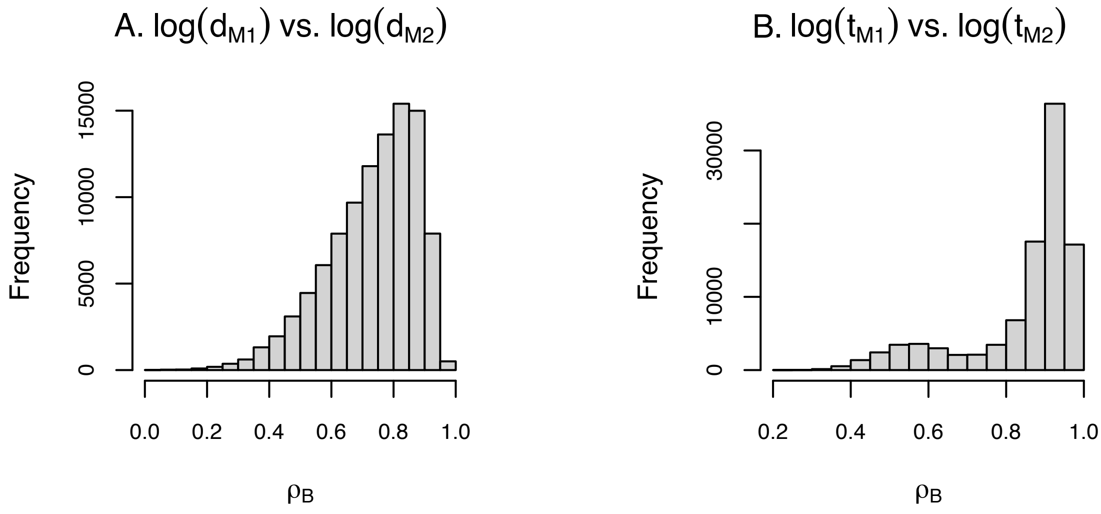


Figure S6: Correlation analyses between the divergence rates ($d$) and substitution times ($t$) calculated by the standard substitution and the population genetics models. To correct the effects of heterogeneous sampling of individuals acrross taxa, the correlation coefficient $\rho_B$ between any two variables was calculated using the bootstrap technique. Specifically, the distribution of $\rho_B$ was obtained by randomly sampling one variable value per taxon at each draw. We employed 1000 bootstraps.

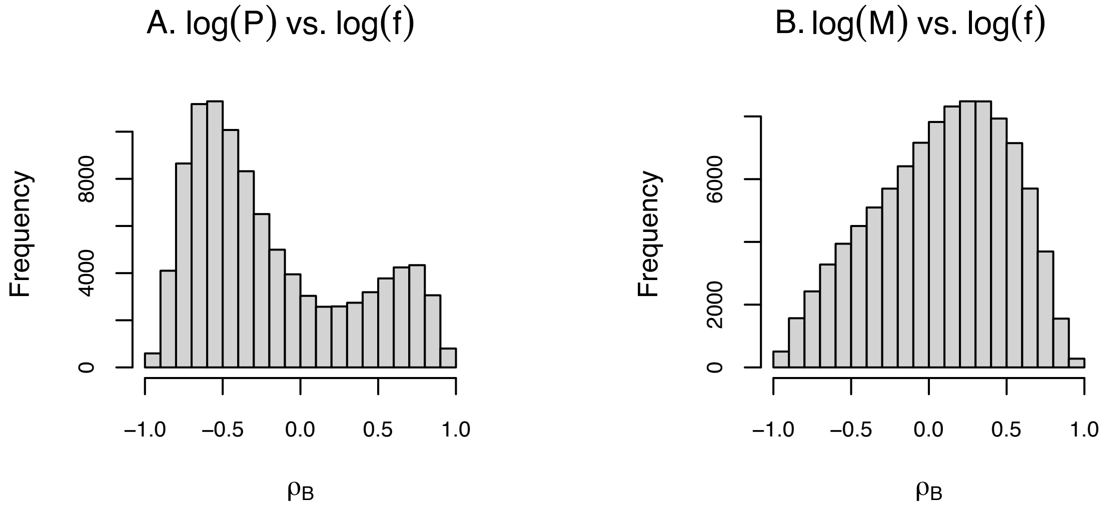


Figure S7: Correlation analyses between the the excess of fixed sites ($f$) and the proportion of polymorphic sites ($P$) and number of sampled individuals ($M$; for diplois, $M$ represents the number of haploid individuals). To correct the effects of heterogeneous sampling of individuals acrross taxa, the correlation coefficient $\rho_B$ between any two variables was calculated using the bootstrap technique. Specifically, the distribution of $\rho_B$ was obtained by randomly sampling one variable value per taxon at each draw; we employed 1000 bootstraps.

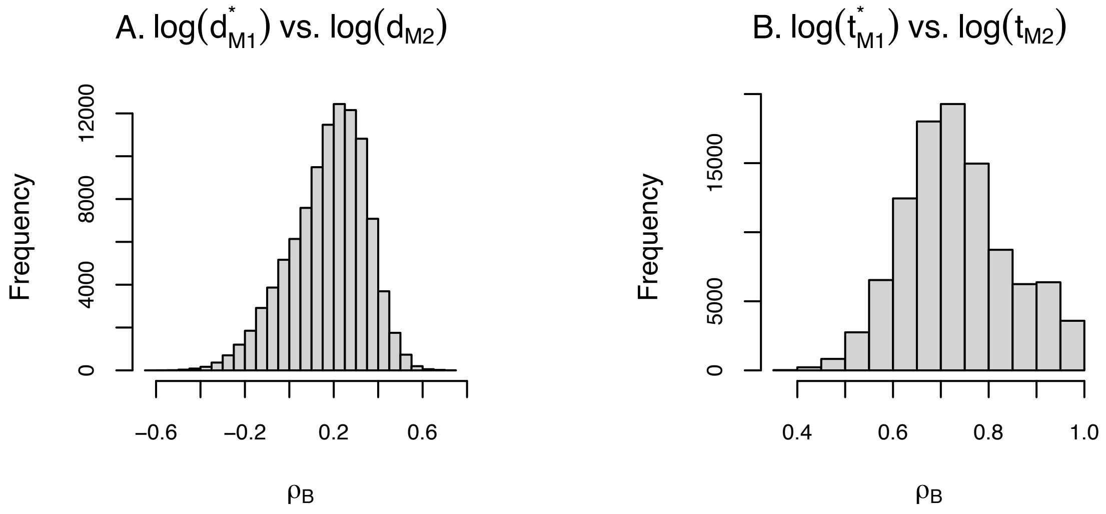


Figure S8: Correlation analyses between the population-informed and naïve estimates of the divergence rate (respectively $d$ and $d^*$) and substitution times (respectively $t$ and $t^*$). To correct the effects of heterogeneous sampling of individuals acrross taxa, the correlation coefficient $\rho_B$ between any two variables was calculated using the bootstrap technique. Specifically, the distribution of $\rho_B$ was obtained by randomly sampling one variable value per taxon at each draw; we employed 1000 bootstraps.
